# Molecular basis of the autoregulatory mechanism of motor neuron-related splicing factor 30

**DOI:** 10.1101/2025.03.04.641417

**Authors:** Keiichi Izumikawa, Tatsuya Shida, Yuuka Onodera, Yuito Tashima, Sotaro Miyao, Tomomi Suda, Yasuyuki Suda, Minoru Sugihara, Tamotsu Noguchi, Masami Nagahama

## Abstract

Motor neuron-related splicing factor 30 (SPF30, also known as SMNDC1) is a paralog of the survival motor neuron protein that regulates the expression of various genes by affecting mRNA splicing. SPF30 has an autoregulatory mechanism that controls its expression. However, the detailed molecular mechanisms determining cellular levels of SPF30 remain unclear. Here, we demonstrated that SPF30 expression was controlled via the negative autoregulatory feedback, whereby increased SPF30 expression caused the inclusion of cassette exon within intron 2 and/or the generation of a newly spliced variant with exon 4a (produced by splicing 17 bp upstream of the canonical intron 3 and exon 4 junctions). Altered transcripts with cassette exon or exon 4a were subjected to nonsense-mediated mRNA decay, leading to reduced SPF30 mRNA levels. Conversely, the loss of SPF30 protein resulted in a drastic reduction in exon 4a inclusion compared to cassette exon inclusion, suggesting that exon 4a inclusion contributes more to adjusting SPF30 expression levels. An *in vivo* splicing assay designed to reflect exon 4a inclusion levels demonstrated that a short stretch of sequence within exon 4 of SPF30 mRNA was required for exon 4a inclusion. Additionally, the C-terminal region of SPF30 was crucial for the autoregulatory mechanism. Specifically, the C-terminal region of SPF30, including the latter part of α-helix and a kink-like structure, was required for binding to RNA containing exon 4a. Collectively, these results reveal the molecular basis of the autoregulatory mechanism underlying SPF30 gene expression.

## Introduction

Gene expression is regulated by diverse mechanisms, including alternative splicing, which provides the cellular homeostasis required for maintaining developmental processes or diverse tissue specificities (1, 2). Dynamic variations in splicing factors or RNA-binding proteins that access pre-mRNAs help determine the fate of cells and tissues (3, 4). Certain genes involved in splicing events can modify their own splicing variations to regulate their expression levels, a mechanism known as autoregulation (5). A common characteristic of autoregulated genes is that their cellular expression levels must be tightly controlled, as substantial changes in their expression can severely impact cellular and organismal homeostasis, potentially threatening survival. For example, TAR DNA-binding protein 43 (TDP-43), a product of the TARDBP gene, is controlled by an autoregulatory mechanism, and its absence results in embryonic lethality (6–8). Elevated TDP-43 expression, as observed in amyotrophic lateral sclerosis (ALS), reduces cell proliferation and causes cell death. In diseases such as ALS, motor neuron death occurs due to cytoplasmic aggregation (9–11). Genes involved in splicing events, such as SF2ASF, serine/arginine-rich splicing factor 2 (SC35), PTB, hnRNP L, hnRNP D, and Chtop, often have autoregulatory mechanisms (12–17). The factors involved in splicing events serve critical roles in maintaining cellular homeostasis by maintaining constant protein levels.

Splicing factor 30 (SPF30), also known as survival motor neuron domain-containing protein 1 (SMNDC1), is a paralog of the survival motor neuron (SMN) protein and is characterized as a dimethylarginine-binding protein with a Tudor domain (18–20). SPF30 is an essential splicing factor required for the assembly of three small nuclear ribonucleoproteins (snRNPs)—namely, U4, U5, and U6— into the spliceosome (21). During this process, SPF30 binds to the 17S U2 snRNP, promoting recruitment of the U4, U5, and U6 snRNPs to the pre-spliceosome (21–23). Although SPF30 is considered an essential splicing factor, changes in SPF30 expression can alter the splicing patterns (particularly intron retention and exon skipping) of specific gene sets containing chromatin-remodeling factors, such as Atrx, in murine cells (24, 25). These results suggest that SPF30 acts as a splicing factor that regulates tissue-specific gene expression rather than as an essential splicing factor. In addition to its functional association with splicing-related proteins, SPF30 can regulate gene expression by binding to chromatin-remodeling factors or components of RNA exosomes containing MTR4, resulting in altered cellular chromatin accessibility or RNA metabolism (25, 26).

Despite the important cellular functions of SPF30, the regulatory mechanisms underlying its expression levels remain unclear. Some investigators have predicted that SPF30 may possess an autoregulatory mechanism, controlling its own splicing events—particularly by including a cassette exon within intron 2 of SPF30 and promoting mRNA degradation via nonsense-mediated mRNA decay (NMD) (24, 27, 28). Despite accumulating evidence that SPF30 expression causes exon inclusion within intron 2, the mechanism whereby SPF30 regulates SPF30 expression remains unknown. Here, we show that, in addition to promoting the inclusion of the cassette exon into intron 2, SPF30 negatively controls its own protein expression by affecting the splicing of its pre-mRNA after binding near the intron 3/exon 4 junction. In particular, exon 4a, which is produced by splicing 17 bp upstream of the junction between the canonical intron 3 and exon 4 (rather than cassette exon inclusion), contributed more to the SPF30 autoregulatory mechanism. This activity, in turn, facilitated the generation of an alternative variant of exon 4a-retained SPF30 mRNA, which was subsequently degraded via NMD.

## Results

### Exogenous SPF30 expression reduces endogenous SPF30 protein and mRNA levels

We established an Flp-In T-REx 293 (293FTR) cell line capable of inducing the expression of a single copy of a C-terminally FLAG-tagged and 6xHis-tagged variant of SPF30 (SPF30-FLAG) transgene. Western blot analysis showed that SPF30-FLAG expression plateaued at 48 h after induction with doxycycline (Dox) (Fig. 1*A*). Immunocytochemical staining showed that SPF30-FLAG was localized to nuclear speckles, partly coinciding with the speckle marker SC35, similar to endogenous SPF30, indicating that SPF30-FLAG localized similarly to endogenous SPF30 (Fig. 1*B* and Fig. S1) (29). Western blot analysis of SPF30-FLAG and endogenous SPF30 showed that SPF30-FLAG had a higher molecular weight than endogenous SPF30, which was attributable to the FLAG and 6xHis protein sequences, enabling the distinction of both proteins (Fig. 1*A*). SPF30-FLAG expression reduced endogenous SPF30 expression by up to 60% after 48 h when compared with that in uninduced cells, whereas Dox treatment did not affect endogenous SPF30 expression in parental 293FTR cells (Fig. 1*A*). Quantitative RT-PCR (RT-qPCR) analysis confirmed that endogenous SPF30 mRNA expression, as shown by the 3’UTR primer set, decreased by up to 60% after SPF30-FLAG expression for 48 h compared with that in uninduced cells (Fig. 1, *C* and *D*). In contrast, Dox induction did not affect endogenous SPF30 mRNA expression in 293FTR cells (Fig. 1*D*). Therefore, downregulation of endogenous SPF30 protein expression due to SPF30-FLAG expression correlated with decreased endogenous SPF30 mRNA expression (Fig. 1, *A* and *D*). These results suggest that the observed reduction in endogenous SPF30 protein levels upon SPF30-FLAG expression was caused at the mRNA level by a negative-feedback mechanism involving SPF30 gene expression.

**Figure 1.**
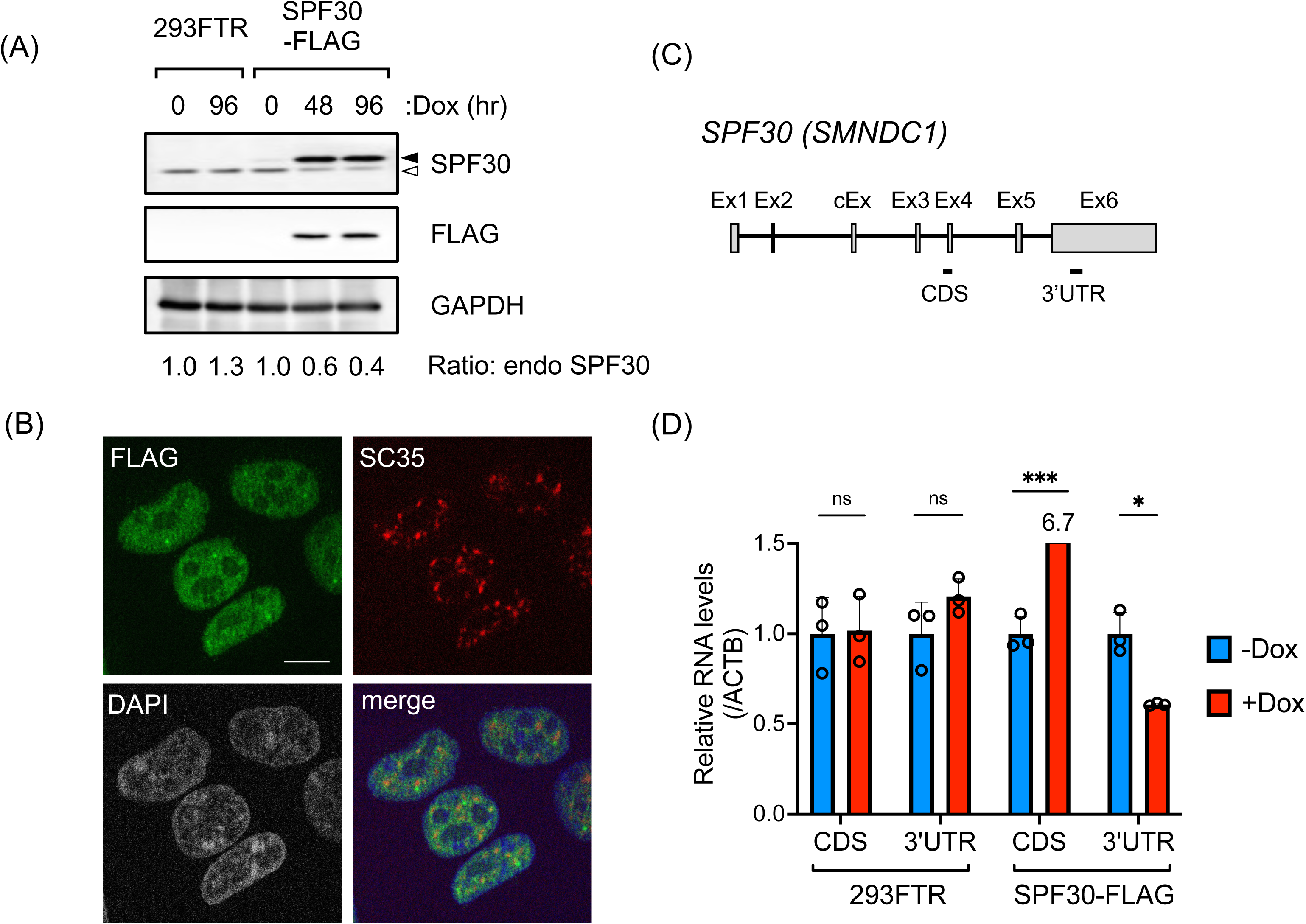
Endogenous SPF30 mRNA is regulated by SPF30 protein expression levels. (*A*) Endogenous SPF30 or SPF30-FLAG expression levels were analyzed by western blotting using cells capable of SPF30-FLAG induction and parental (control) 293FTR cells following Dox treatment for the indicated durations. The closed and open arrowheads indicate bands corresponding to SPF30-FLAG and endogenous SPF30, respectively. The antibodies used are indicated on the right side of each blot. The relative band intensities of endogenous SPF30 to uninduced cells (0 h Dox treatment) are shown. The data presented represent the averages of three independent experiments. (*B*) Immunofluorescent images of SPF30-FLAG and SC35 following SPF30-FLAG induction. Using SPF30-FLAG-induced cells treated with 1 µg/ml Dox for 24 h, SPF30-FLAG and SC35 were stained with primary anti-FLAG and anti-SC35 antibodies and then stained with secondary Alexa488-conjugated anti-rabbit IgG and Alexa595-conjugated anti-mouse IgG. SC35 was detected as a nuclear speckle marker. Scale bar: 10 µm. (*C*) Schematic representation of the SPF30 gene. Exons (Ex; open boxes) and introns (lines) of SPF30 are shown. DNA fragments of the coding sequence (CDS) and 3’-untranslated region (UTR) amplified by RT-qPCR are shown. cEx; cassette exon. (*D*) RT-qPCR analysis of exogenous and endogenous SPF30 mRNA expression. SPF30-FLAG-inducible cells or 293FTR cells were treated with Dox for 48 h, and endogenous SPF30 mRNA expression was analyzed by RT-qPCR using a primer set targeting the 3’-UTR. Total (endogenous and exogenous) SPF30 mRNA levels were measured using a CDS primer set. Data are presented as mean ± standard deviation (SD) of the three independent experiments. *P < 0.05, ***P < 0.001, ns; not significant (Welch’s test).

### SPF30 overexpression promotes the inclusion of exon 4a, generating transcripts with premature terminal codon

SPF30 expression is regulated by the inclusion of a cassette exon within intron 2 via alternative splicing, generating transcripts with premature terminal codons (PTCs) within exon 3 that are degraded via NMD, which is thought to affect SPF30 expression (24, 28). To define the degradation pathway of the alternatively spliced SPF30 transcripts in the autoregulatory process, we examined the effect of cycloheximide (CHX; an NMD inhibitor) on transcripts degraded by NMD with or without SPF30-FLAG induction. RT-PCR conducted using the Ex1/Ex6 and Ex1/Ex4 primer sets revealed increased abundances of aberrant transcripts approximately 1000 bp (Ex1/Ex6) and 700 bp (Ex1/Ex4) long upon CHX treatment. These aberrant transcripts were larger than the PCR products from mature transcripts by 800 bp (Ex1/Ex6) or 500 bp (Ex1/Ex4), respectively, suggesting that they contained a cassette exon within intron 2 (Fig. 2, *A* and *B*) (24, 28).

**Figure 2.**
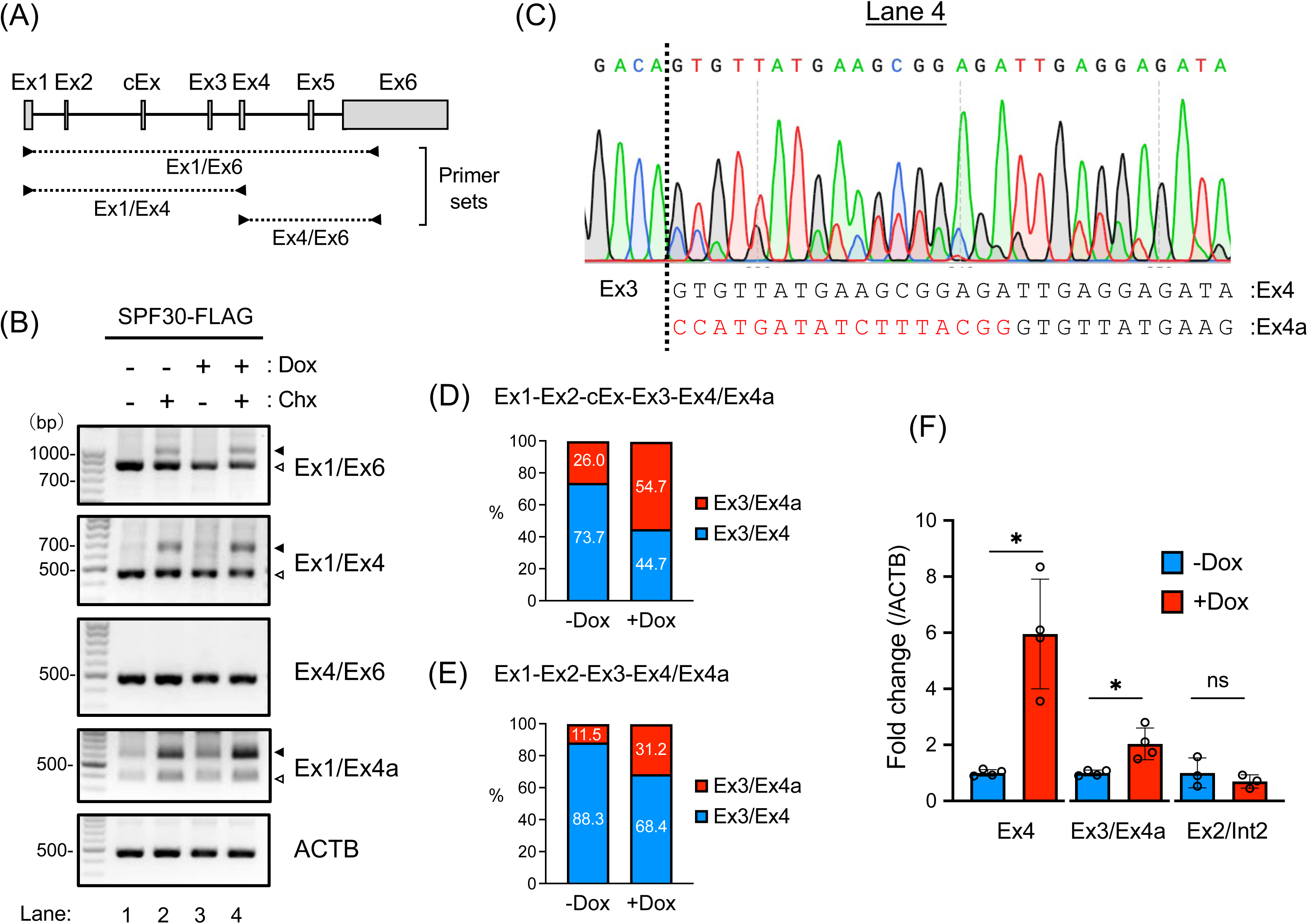
SPF30 overexpression causes inclusion of the cassette exon and/or generation of exon 4a instead of exon 4. (*A*) Schematic representation of the SPF30 gene, showing exons (Ex; open boxes) and introns (lines). Primer sets used to detect SPF30 mRNA in the experiments represented in panel *B* are shown. cEx; cassette exon. (*B*) SPF30-FLAG cells were treated with Dox for 24 h and/or CHX for 4 h, and endogenous SPF30 mRNA variants were analyzed by RT-PCR. The primer sets used are shown at the right of each gel image. The open and closed arrowheads show the transcripts for exons 1 to 4, and for exons 1 to 4 containing the cassette exon, respectively. ACTB was detected as a control. (*C*) Sanger sequencing was performed with a DNA fragment amplified using the Ex1/Ex4 primer set. The open arrowhead in lane 4 of panel *B* indicates the region analyzed. The DNA sequence was annotated and described based on the sequencing chromatogram. (*D*, *E*) SPF30-FLAG cells treated with (+Dox) or without (−Dox) Dox for 24 h were treated with CHX for 2 h, and SPF30 mRNA was amplified using the Ex1/Ex4 primer set. The amplified DNA fragments, with (*D*) or without (*E*) the cassette exon, were analyzed by deep sequencing. The graphs show the percentages of DNA fragments containing exons 3 and 4 (Ex3/Ex4) and exons 3 and 4a (Ex3/Ex4a). (*F*) The relative expression levels of SPF30 transcripts containing exon 4a or the cassette exon in SPF30-FLAG uninduced (−Dox) or induced (+Dox) cells were measured by RT-qPCR. Data are presented as the mean ± SD of the three or four independent experiments. *P < 0.05, ns; not significant (Welch’s test).

Next, deep and Sanger sequencing were performed to annotate the sequences of the SPF30 transcripts with cassette exons and mature SPF30 transcripts. The cassette exon sequence was annotated to bp 4214–4403 of the SPF30 gene (NG_008284) by deep sequencing of the PCR-amplified DNA fragments (Fig. S2, *A* and *B*). Sanger sequencing of mature SPF30 transcripts (500 bp) following SPF30-FLAG induction and CHX treatment revealed that, in addition to mature transcripts containing exons 1–4, splice variants consisting of exons 1–3 and a longer exon 4 were present in the 500 bp PCR-amplified band obtained using the Ex1/Ex4 primers (Fig. 2*B*, lane 4). The longer version of exon 4 was identified as a variant produced by splicing 17 bp upstream of the canonical junction of introns 3 and 4 and was named as exon 4a (Fig. 2*C*).

In contrast, only mature transcripts were detected in untreated cells, whereas faint transcript signals of exon 4a were detected in mature transcripts after CHX treatment without SPF30-FLAG induction (Fig. S2, *C*–*E*). These results indicated that SPF30-FLAG overexpression generated splice variants with exon 4a containing a PTC within exon 4, which was therefore inferred to be degraded by NMD. Deep sequencing of transcripts with or without the cassette exon upon CHX treatment also revealed that SPF30 overexpression increased the percentage of exon 4a inclusion from 26.0% to 54.7% in transcripts with the cassette exon and from 11.5% to 31.2% in transcripts without the cassette exon (Fig. 2, *D* and *E*). RT-PCR performed using a specific primer for exon 4a revealed that SPF30 transcripts containing exon 4a were more abundant after CHX treatment (Fig. 2*B*, see the data presented for Ex1/Ex4a). These results indicate that the expression of transcripts with exon 4a increased by SPF30-FLAG expression but were degraded by NMD (Fig. 2*B*, see the data presented for Ex1/Ex4a).

RT-qPCR using specific primers for exon 3 and exon 4a (Ex3/Ex4a), which were designed to detect all transcripts with exon 4a, also revealed that SPF30-FLAG expression significantly increased the total exon 4a inclusion levels of endogenous SPF30 mRNA, whereas the abundance of transcripts with the cassette exon (detected using a primer for exon 2 and intron 2 [Ex2/Int2]) did not change (Fig. 2*F*). These results indicate that increased SPF30 expression led to exon 4a inclusion, resulting in SPF30 transcript degradation via NMD, which underlies the autoregulatory mechanism of SPF30 gene expression.

### Reduction of SPF30 protein expression results in the accumulation of mature SPF30 mRNA by inhibiting the inclusion of cassette exon and exon 4a

To test the autoregulatory mechanism of SPF30, we used the auxin-inducible degron (AID) 2 protein-knockdown system to examine splice-variant levels of SPF30 upon SPF30 protein depletion. We established a single clonal SPF30-mAID-mCherry cell line (SPF30-mAID cells), in which the sequence encoding the minimal functional AID and mCherry tandem tag was inserted into the 3’ end of the protein-coding sequence of the endogenous SPF30 gene (SPF30-mAC) in both alleles of the HCT116 cell line. This cell line stably expressed the F74G variant of the *Arabidopsis thaliana* TIR1 protein, i.e., OsTIR1 (F74G) (Fig. 3*A*) (30).

**Figure 3.**
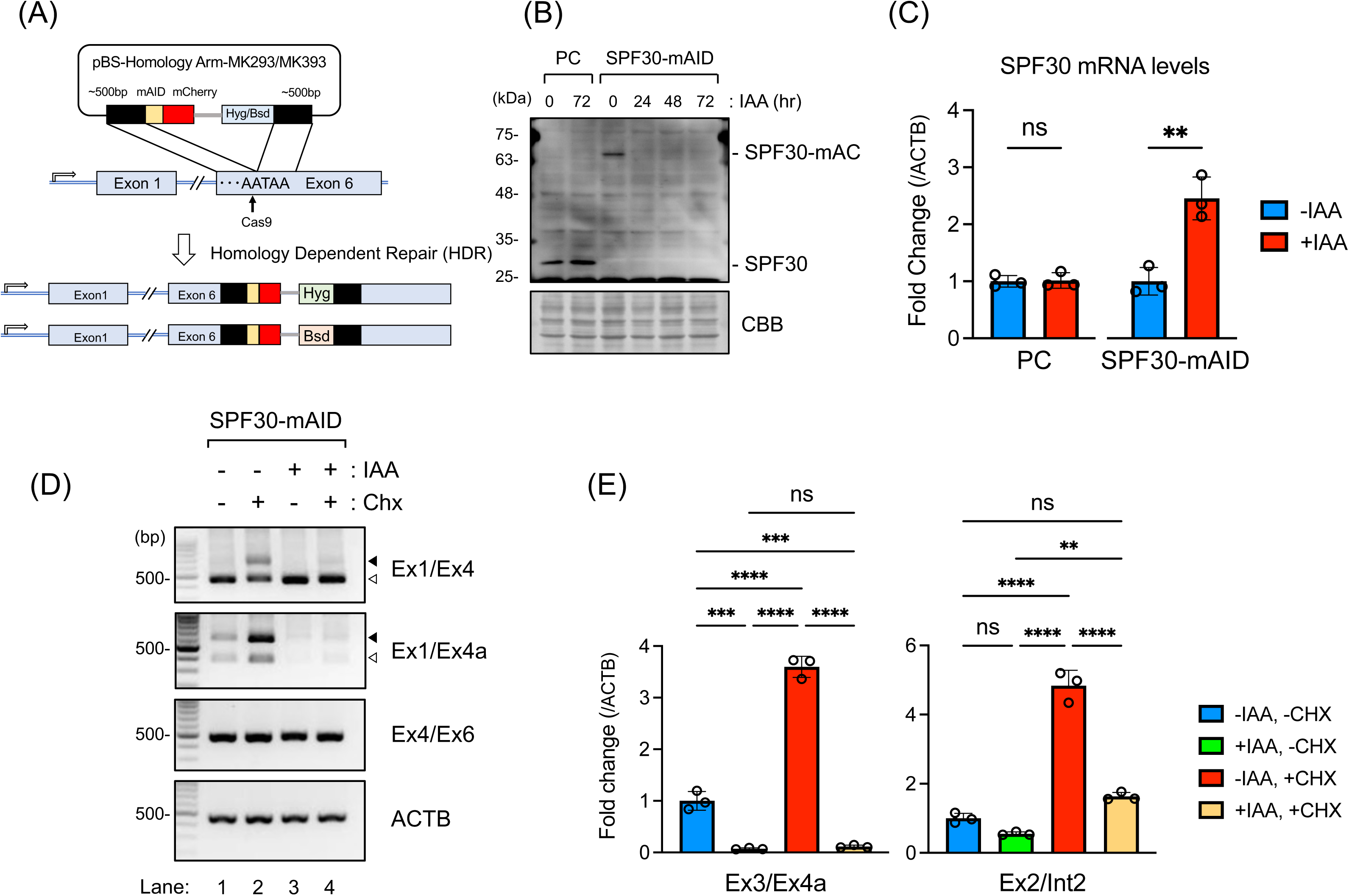
SPF30 knockdown and splice variant analysis in SPF30-mAID cells using the AID2 system. (*A*) Schematic representation of the establishment of an SPF30 protein-knockdown cell line (SPF30-mAID) that expressed SPF30-mAID-mCherry, using the AID2 system. (*B*) SPF30-mAID cells were treated with 0.25 µM IAA for the indicated durations, and SPF30 expression was detected via western blot analysis with an anti-SPF30 antibody. Parental HCT116 cells (PC) were used as a control. Total amounts of the loaded proteins were detected by CBB staining. (*C*) The relative expression levels of SPF30 transcripts in SPF30-mAID cells treated with IAA (+IAA) or not treated (−IAA) were measured via RT-qPCR analysis. A primer set targeting exon 4 was used for detection. PCs were used as a control. Data are presented as the mean ± SD of three independent experiments. **P < 0.01, ns; not significant (unpaired t-test). (*D*) SPF30-mAID cells were treated with 0.25 µM IAA for 72 h and/or CHX for 2 h, and SPF30 transcripts containing exon 4a or the cassette exon were detected by RT-PCR. The primer sets used are indicated at the right of each gel image. The open and closed arrowheads show the transcripts for exons 1 to 4, and for exons 1 to 4 containing the cassette exon, respectively. ACTB was detected as a control. (*E*) The relative expression levels of SPF30 transcripts containing exon 4a (Ex3/Ex4a) or the cassette exon (Ex2/Int2) in SPF30-mAID cells were measured via RT-qPCR analysis. SPF30-mAID cells were treated with IAA (+IAA) or were not treated (−IAA) for 72 h, followed by treatment with or without CHX for 2 h. Data are presented as the mean ± SD of the three independent experiments. **P < 0.01, ***P < 0.001, ****P < 0.0001, ns; not significant (Tukey’s test).

In SPF30-mAID cells, SPF30-mAC expression decreased steadily, even after 24 h of 5-phenyl-1H-indole-3-acetic acid (IAA) treatment (Fig. 3*B*). Using this cell line, we examined SPF30 transcript levels after SPF30 protein knockdown. RT-qPCR analysis revealed that total SPF30 mRNA significantly (P = 0.0048) accumulated by approximately 2-fold upon SPF30 protein knockdown. In contrast, SPF30 mRNA levels did not change in the parental HCT116 cells (Fig. 3*C*). To detect transcripts degraded by NMD, SPF30-mAID cells were treated with CHX. RT-PCR analysis revealed increased inclusion of the cassette exon and exon 4a upon CHX treatment without SPF30 protein knockdown in SPF30-mAID cells, as also observed in 293FTR cells (Fig. 2*B*; Fig. 3*D*, lane 2). In contrast, the inclusion of the cassette exon and exon 4a decreased markedly upon SPF30 knockdown in CHX-treated cells, as shown by PCR using the primer sets Ex1/Ex4 and Ex1/Ex4a, respectively (Fig. 3*D*, lanes 3 and 4).

RT-qPCR analysis of exon 4a showed that SPF30 knockdown decreased the total transcript levels of exon 4a to as low as one-fourteenth of SPF30 transcript levels in cells without IAA and CHX treatment, and one-thirtieth in cells with IAA and CHX treatment. Additionally, CHX treatment did not affect exon 4a levels upon SPF30 knockdown (Fig. 3*E*, left). In contrast, transcripts containing the cassette exon, detected with the Ex2/Int2 primer set, were reduced by 45% without CHX and by 66% with CHX following SPF30 knockdown (Fig. 3*E*, right). These results indicate that SPF30 functioned as a splicing factor, leading to the inclusion of cassette exon and exon 4a. In particular, SPF30 played a critical role in the inclusion of exon 4a, when compared with the inclusion of the cassette exon, and controlled its own expression levels through the autoregulatory mechanism of SPF30 by controlling exon 4a inclusion.

### An element within exon 4 is required for exon 4a inclusion in SPF30 autoregulation

To determine the mRNA sequences required for the exon 4a inclusion, an *in vivo* splicing assay was performed using NanoLuc Luciferase (Nluc). For this purpose, we constructed a minigene to detect exon 4a inclusions, where the DNA region from exons 3 and 4 of SPF30 was inserted upstream of the Nluc gene, followed by a sequence rich in proline, glutamic acid, serine, and threonine (PEST sequence) that was targeted for degradation as a short-lived protein (Fig. 4, *A* and *B*). In the splicing assay, transcripts containing exons 3 and 4 were translated as Nluc-containing proteins, whereas the inclusion of exon 4a caused a frameshift in the Nluc-coding sequence, resulting in the loss of the Nluc signal. The minigene was transfected into SPF30-mAID cells, and exon 4a inclusions were detected by measuring Nluc signals with or without SPF30 knockdown. SPF30 knockdown following IAA treatment resulted in higher Nluc activity compared with that in the untreated group, suggesting that transcripts with exon 4 were upregulated due to inhibited exon 4a inclusion caused by SPF30 depletion (Fig. 4*C*, WT).

**Figure 4.**
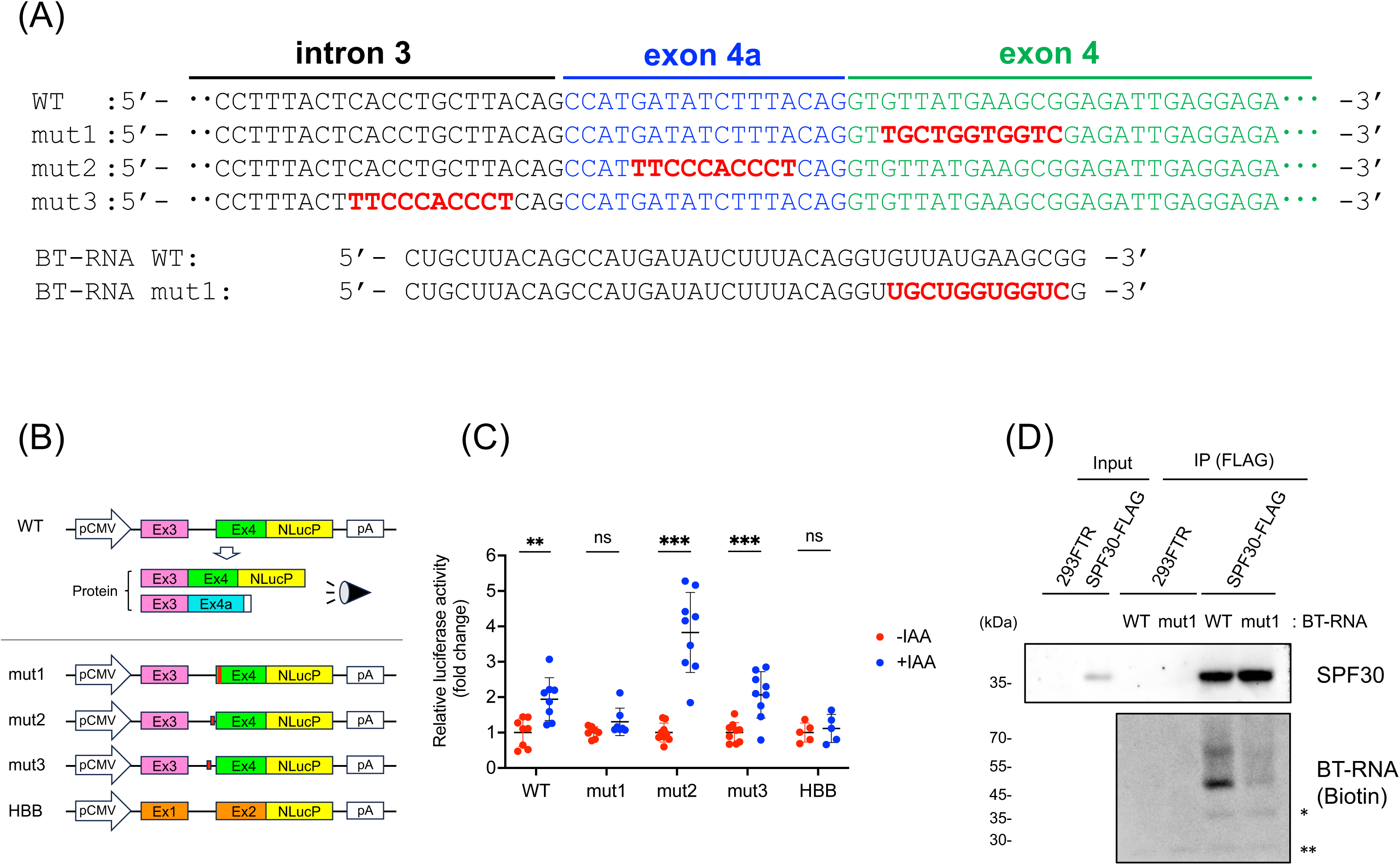
RNA sequences within exon 4 are required for exon 4a inclusion through the autoregulatory mechanism of SPF30. (*A*) DNA sequences of WT, mut1, mut2, and mut3 around intron 3, exon 4a, and exon 4 in an Nluc-expressing plasmid are shown. The sequences of BT-RNA WT and BT-RNA mut1 are shown. (*B*) Schematic representation of the minigenes used in an *in vivo* splicing assay. The DNA sequence from exons 3–4 of the SPF30 gene was inserted upstream of Nluc plasmid DNA. The transcripts with exons 3 and 4 were translated into a protein containing the Nluc protein sequence. DNA elements within exon 4 (mut1), exon 4a (mut2), or intron 3 (mut3) were replaced with the corresponding sequences of the human HBB gene. (*C*) The relative luciferase activities of HCT116 cells transfected separately with each NlucP plasmid were analyzed in *in vivo* splicing assays. Data are presented as the mean ± SD of five to nine independent experiments. **P < 0.01, ***P < 0.001, ns; not significant (Welch’s test). (*D*) Direct binding of SPF30 to exon 4a-containing RNA was analyzed with a UV cross-linking-based method with immunoprecipitated SPF30-FLAG and BT-RNA WT containing the unique sequence of exon 4a. Another BT-RNA, in which the RNA sequence within exon 4 was replaced with that of HBB, was used as a negative control (mut1). BT-RNA was detected with streptavidin-conjugated HRP.

Next, three additional minigenes (mut1–3) and a control human hemoglobin subunit beta (HBB) minigene were constructed (Fig. 4, *A* and *B*). In the modified minigenes, the DNA sequence in the vicinity of exon 4a was replaced by the HBB minigene sequence (NG_059281.1). Specifically, in mut1, the region immediately downstream of the intron 3–exon 4 junction of SPF30 was replaced by the sequence from exon 2 of the HBB minigene (NG_059281.1, 275–286). In mut2, the sequence within exon 4a of SPF30 was replaced by intron 1 of the HBB minigene (NG_059281.1, 260–269). In mut3, the region upstream of the intron 3–exon 4a junction of SPF30 was replaced by intron 1 of the HBB minigene. As a control HBB minigene, exons 1–2 of HBB were inserted upstream of Nluc.

In the *in vivo* splicing assays performed using the unmodified HBB minigene, SPF30 knockdown did not change the Nluc signals, indicating that a splicing-inhibitory sequence was not present in the HBB minigene (Fig. 4*C*, HBB). In splicing assays with the modified minigenes, mut1 did not affect Nluc signals upon SPF30 knockdown, whereas mut2 and mut3 increased Nluc signals similar to the WT minigene. These results indicate that the sequence corresponding to the region from the third to the thirteenth nucleotide of exon 4 was required for regulating exon 4a inclusion based on SPF30 protein expression, which provides support to the autoregulatory mechanism of SPF30 (Fig. 4*C*). We next assessed the possibility that the exon 4 region responsible for splicing regulation was necessary for SPF30 protein binding to SPF30 mRNA. In an *in vitro* protein–RNA binding assay, SPF30-FLAG inducible cells were incubated with synthetic biotinylated RNA oligonucleotides (BT-RNA) derived from wild-type (WT) or mut1 (Fig. 4, *A* and *D*; Fig. S3). As shown by the detection of the BT-RNA band at approximately 45 kDa, SPF30-FLAG directly bound to BT-RNA WT but not to BT-RNA mut1, indicating that the sequence within exon 4 was required for SPF30 binding (Fig. 4*D*). An upper band of approximately 70 kDa in the SPF30–BT-RNA complex was also detected, suggesting that another unidentified protein component, consisting of SPF30 and BT-RNA, may exist in the complex (Fig. 4*D*).

### The C-terminal region of SPF30 is responsible for its autoregulation

SPF30 is composed of a tandem α-helical N-terminal domain, a Tudor domain, and a C-terminal domain with an α-helix and intrinsically disordered region (19, 22, 29). AlphaFold2 predictions generated five structural models of the SPF30 structure (31) . Among them, only the fourth model of SPF30 exhibited potential binding with the 64-nucleotide (nt) RNA region containing exon 4a; therefore, we adopted the fourth model as an SPF30 structural model (Fig. S4*A*). To determine the protein region in SPF30 responsible for its autoregulation through the inclusion of the cassette exon and exon 4a of SPF30, we established five 293FTR cell lines capable of inducibly expressing deletion mutants (ΔN, ΔTudor, ΔC1, ΔC4, and ΔC5) of SPF30 (Fig. 5*A*; Fig. S4*B*). Using these cell lines, we examined endogenous SPF30 mRNA levels upon expression of each deletion mutant, using RT-qPCR (Fig. 5*B*). ΔN and ΔTudor expression downregulated endogenous SPF30 expression, whereas the ΔC1, ΔC4, and ΔC5 mutants did not affect endogenous SPF30 levels, indicating that the C-terminal region of SPF30 participated in autoregulating SPF30 expression (Fig. 5*B*).

**Figure 5.**
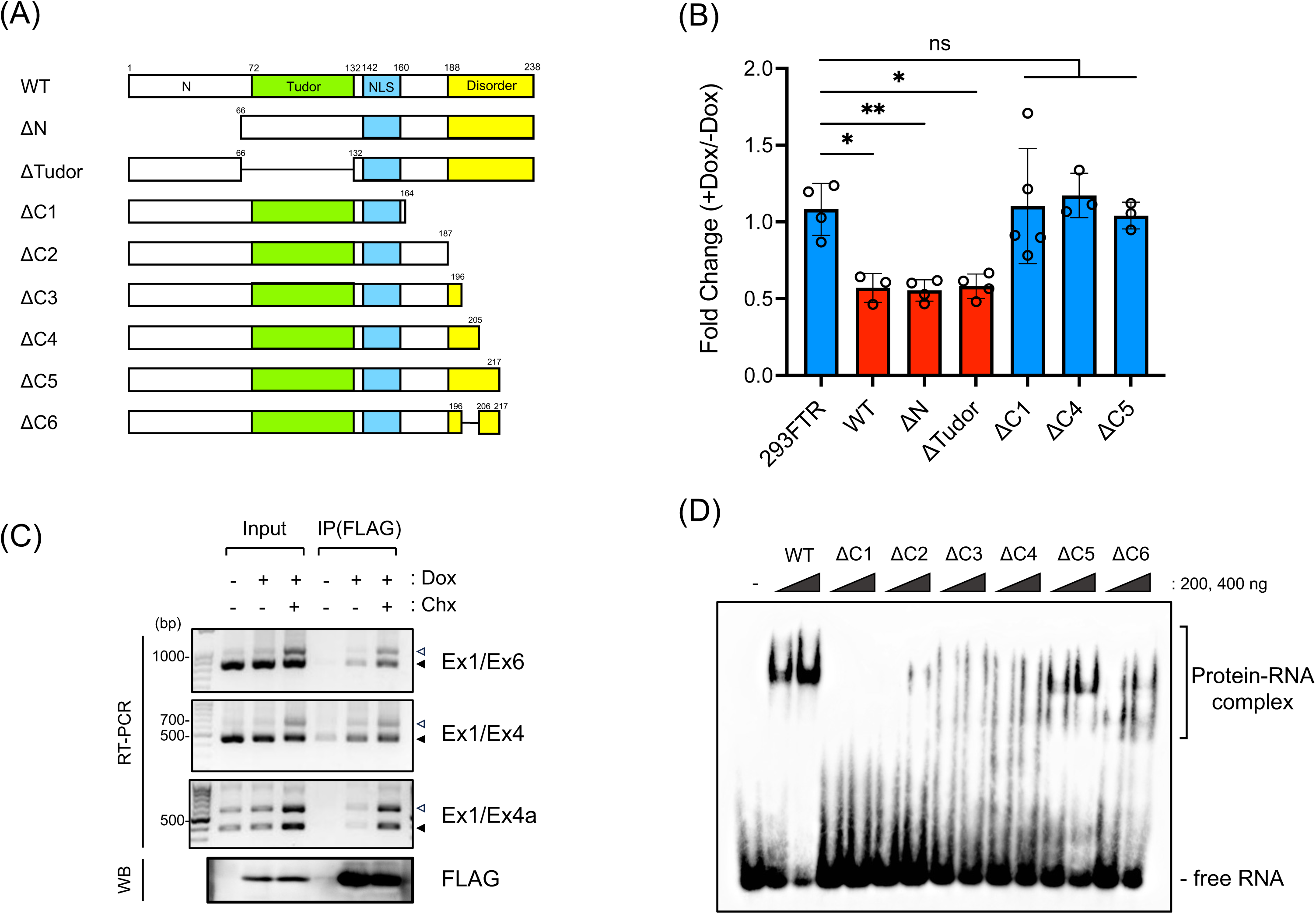
C-terminal elements of SPF30 are required for the autoregulatory mechanism of SPF30. (*A*) Schematic representation of SPF30-FLAG mutants. (*B*) Endogenous SPF30 mRNA levels in SPF30-FLAG mutant-inducible cells were detected via RT-qPCR analysis using a primer set targeting exon 4. 293FTR cells and WT SPF30-FLAG (WT) were used as negative and positive controls, respectively. Data are presented as the mean ± SD of the three to four independent experiments. *P < 0.05, **P < 0.01, ns; not significant (Dunnett’s test). (*C*) SPF30-FLAG-inducible cells treated with or without Dox, as well as CHX, were immunoprecipitated using an anti-FLAG antibody, and endogenous SPF30 transcripts were detected via RT-PCR with the primer sets indicated at the right of each gel image. SPF30-FLAG protein expression was detected with western blotting with an anti-FLAG antibody. (*D*) EMSAs were performed with BT-RNA and 200 ng or 400 ng of recombinant SPF30 or SPF30 mutants (ΔC1–ΔC6).

Further analyses were performed to determine the SPF30 protein region responsible for RNA binding. RNA immunoprecipitation assays performed using SPF30-FLAG revealed that SPF30 bound SPF30 mRNA containing exon 4a in intact cells (Fig. 5*C*). Given the pivotal role of the C-terminal region of SPF30 in the autoregulatory mechanism of SPF30 gene expression, we evaluated its necessity for binding to RNA containing the unique sequence present in exon 4a. An electrophoretic mobility shift assay (EMSA) was performed using recombinant SPF30 protein and BT-RNA (Fig. S4*C*; WT). EMSA analysis revealed apparent band-shifts of BT-RNA with WT and ΔC5; no band-shift with ΔC1; and slight, smeared band-shifts with ΔC2, ΔC3, and ΔC4 (Fig. 5*D*). The EMSA results suggested that the region of SPF30 containing amino acids (aa) 165–205 can potentially bind with RNA and that the aa 206–217 region or a kink-like structure including aa 197–217 was pivotal for specific binding to RNA containing exon 4a (Fig. S4, *A* and *D*).

EMSA analysis with an additional deletion mutant of SPF30 lacking aa 197–205 and 218–238 (ΔC6) revealed that the binding affinity of ΔC6 to BT-RNA was lower than that of ΔC5, suggesting that sufficient binding of SPF30 to exon 4a-containing RNA requires the kink-like structure in the 197–217 aa region of SPF30 rather than the primary aa sequence of the region (Fig. 5*D*). In EMSAs performed with the aa 197–217 peptide corresponding to the kink-like structure, the peptide did not show any band shift with BT-RNA, indicating that the kink-like structure alone was insufficient for binding to RNA containing exon 4a (unpublished data). Taken together, our RT-qPCR and EMSA results suggested that SPF30 ΔC1 and ΔC4 did not cause autoregulation of SPF30 mRNA due to the loss of efficient SPF30 RNA binding.

Moreover, SPF30 ΔC5 did not cause SPF30 autoregulation due to the loss of other functions unrelated to RNA binding. These findings imply that SPF30 aa 218–238 might participate in an as-yet-unidentified protein binding, which is also involved in the autoregulation mechanism of SPF30.

### The C-terminal α-helix and kink-like structure of SPF30 bound transcripts containing exon 4a

Given that the kink-like structure in the C-terminal region of SPF30 was required for binding to SPF30 transcripts containing exon 4a (Fig. 5*D*), we simulated a docking model between the SPF30 protein and SPF30 mRNA containing the unique sequence of exon 4a using the HDOCK server, a protein– protein/nucleic acid docking web server (http://hdock.phys.hust.edu.cn/) (32, 33). The secondary structure of the pre-mRNA sequence containing intron 3 and exon 4a of SPF30 mRNA was predicted using the Vienna RNAfold web server (34). The 16 nt downstream of the intron 3–exon 4 junction were predicted to form dsRNA structures between exon 4 and 10 nt within a unique 17-bp sequence in exon 4a, with the intron 3-exon 4a junction site predicted to form a loop structure (Fig. S5, *A* and *B*). In the predicted model with the highest score on the HDOCK server, latter half of the C-terminal α-helix and the immediately downstream kink-like structure of SPF30 were bound to the RNA (Fig. 6, *A* and *B*). Notably, the kink-like structure (aa 197–216) of the SPF30 protein seemed to bind the major groove in dsRNA (Fig. 6*B*). The C-terminal α-helix was predicted to be in close proximity with exons 4 and 4a and intron 3 (Fig. S5*C*), and F199, S201, and V209 in the kink-like structure were in close proximity with exon 4a and intron 3 (within 2.5Å) (Fig. 6*C*). In particular, the side chain of F199 and the main chain of V209 were predicted to bind to an nt located at the intron 3–exon 4a junction (Fig. 6*C*). These results support the hypothesis that SPF30 binding to RNA depends on the main chain of aa forming the kink-like structure in SPF30, as well as the latter half of C-terminal α-helix (Fig. 5).

**Figure 6.**
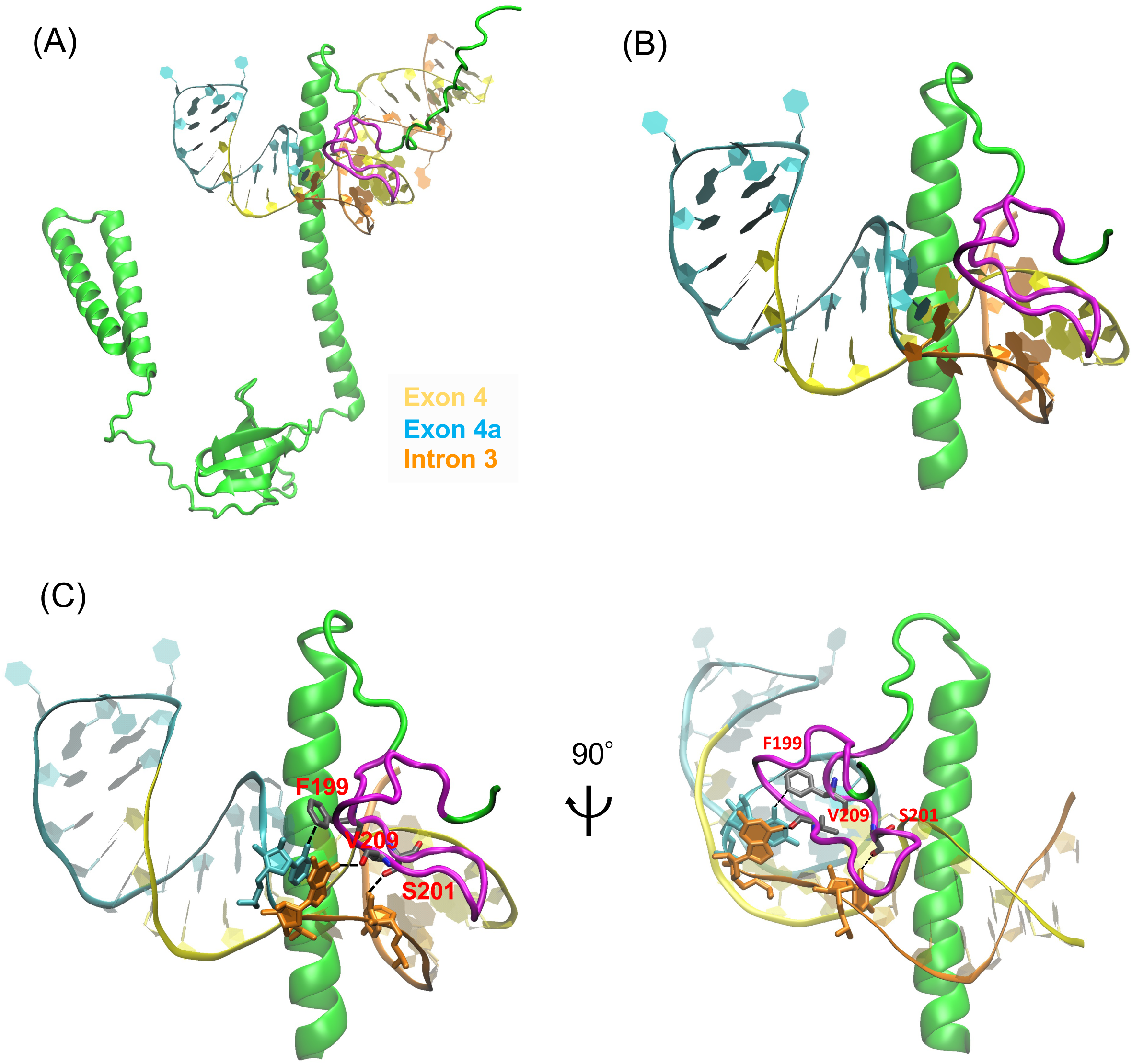
Docking model analysis of SPF30 with RNA containing exon 4a. (*A*–*C*) Molecular docking of SPF30 and RNA containing exon 4a was performed using the HDOCK web server. (*A, B*) The C-terminal α-helix and a kink-like structure (colored magenta) of SPF30 were predicted to be responsible for binding to RNA containing exon 4a. RNA containing exon 4a is shown, with exon 4 in yellow, the unique sequence of exon 4a in cyan, and intron 3 in orange. (*C*) View of the kink-like structure of SPF30 bound to RNA at the intron 3–exon 4a junction.

## Discussion

Here, we demonstrated that cellular SPF30 expression was autoregulated by SPF30 transcripts containing a cassette exon within intron 2 and/or exon 4a, which were alternatively spliced (instead of being processed through the canonical exon 4) and subsequently degraded via NMD (Fig. 7). Furthermore, we showed that SPF30 bound directly to transcripts containing exon 4a with the C-terminal α-helix and the kink-like structure (Figs. 5*D* and 6), resulting in splicing occurring at 17 bp upstream of the intron 3–exon 4 junction (Fig. 2). SPF30 knockdown using the AID2 system, which achieved a higher knockdown efficiency of the SPF30 protein, revealed that cellular SPF30 levels were pivotal for the inclusion of the cassette exon and exon 4a (Fig. 3, *C* and *E*). Thus, a protein-knockdown strategy using the AID2 system is an efficient solution for evaluating the autoregulatory mechanism wherein target genes regulate their mRNA levels.

**Figure 7.**
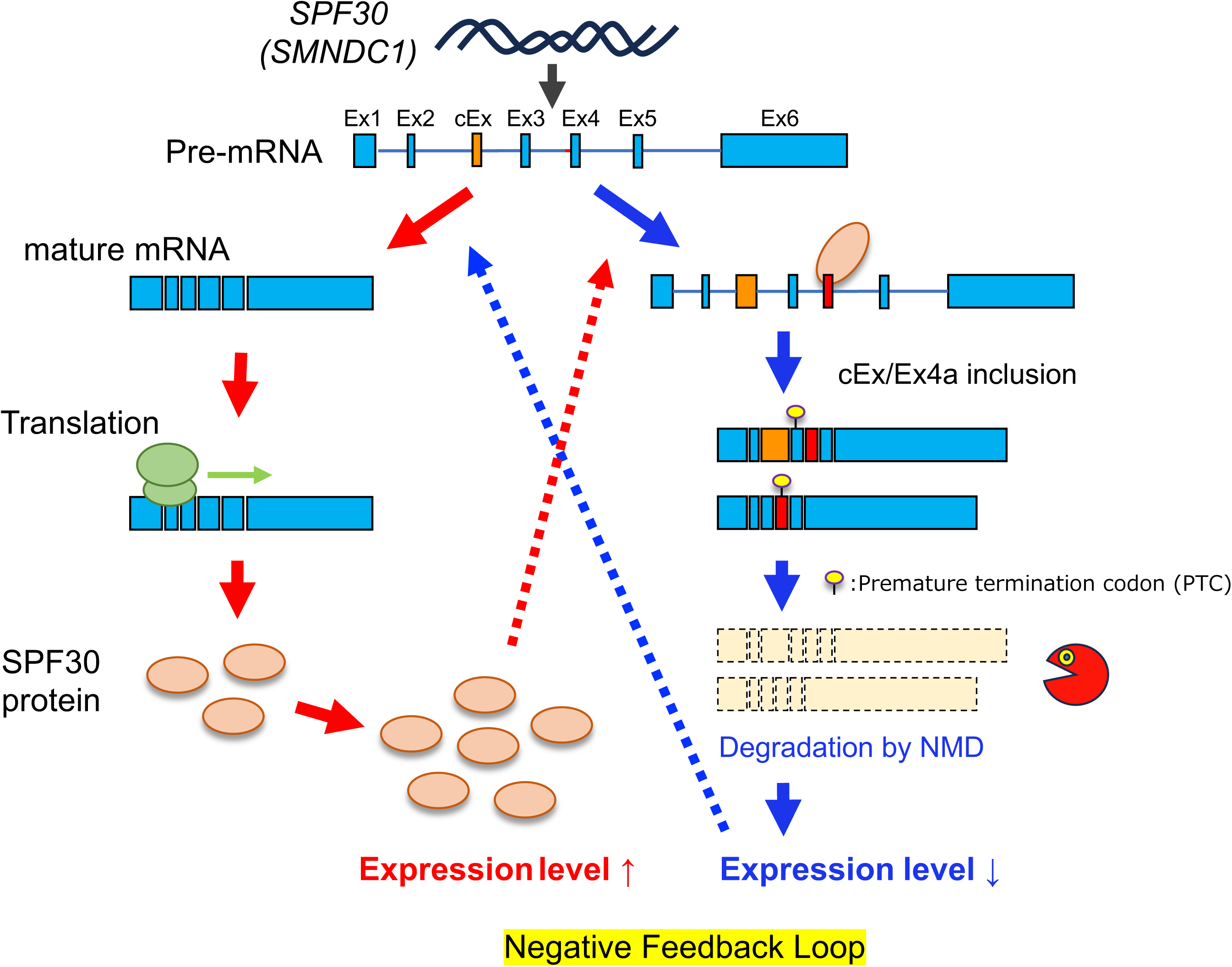
Proposed mechanism whereby SPF30 regulates exon 4a inclusion. SPF30 binds to the element within exon 4 of SPF30 pre-mRNA via its C-terminal α-helix and the kink-like structure and promotes splicing at the intron 3–exon 4a junction, generating transcripts containing exon 4a. Transcripts containing exon 4a generate a PTC, which are degraded via NMD, resulting in reduced SPF30 protein levels.

In this study, we focused on the relative contributions of the cassette exon within intron 2 and exon 4a to the autoregulatory mechanism of SPF30. Deep sequencing of SPF30 transcripts showed that SPF30 overexpression increased the percentage of exon 4a inclusion in transcripts, with and without the cassette exon (Fig. 2, *D* and *E*). Transcripts containing exon 4a were significantly more abundant following SPF30 overexpression, even in the absence of CHX treatment (Fig. 2*F*). These results suggest that the contribution of exon 4a inclusion might be stronger than that of the cassette exon in the autoregulatory mechanism of SPF30. This hypothesis is supported by observations that SPF30 knockdown using the AID2 system reduced the expression of transcripts with the cassette exon by half without CHX treatment and by one-third with CHX treatment. In contrast, transcripts containing exon 4a (with or without CHX treatment) were drastically downregulated, with levels decreasing to as low as one-thirtieth of the original transcript levels (Fig. 3*E*). Taken together, these results indicate that exon 4a responded to changes in SPF30 protein levels more sensitively than the cassette exon, suggesting that exon 4a contributed significantly to SPF30 autoregulation. The Santa Cruz Genome Browser (The University of California) was used to compare the sequence conservation of cassette exon and exon 4a in vertebrates. The cassette exon sequence is conserved among reptiles (e.g., turtles and American alligators), but not among certain fish (e.g., zebrafish and medaka) or birds (e.g., falcons and finches), in which exon 4a sequence is retained (Fig. S6). Therefore, it is predicted that, in the expression of SPF30, an autoregulation mechanism via exon 4a was evolutionarily acquired, followed by an autoregulation mechanism via a cassette exon within intron 2, which developed into the current two-step autoregulatory mechanism in vertebrates.

The detailed mechanism of mRNA splicing involving exon 4a might involve steric hindrance at the junctions of intron 3 and exon 4 or the promotion of splicing recognition at the junctions of intron 3 and exon 4a. Our docking model between the SPF30 protein and SPF30 mRNA containing exon 4a did not provide clear evidence of steric hindrance at the intron 3–exon 4 junction, although the additional effects of interacting proteins with SPF30 were not considered (Fig. 6). In contrast, in the docking model, SPF30 bound to the junctions of introns 3 and 4a, indicating that it might promote splicing (Fig. 6*C*). In addition, knockdown of SPF30 drastically reduced exon 4a splicing, suggesting that SPF30 protein may promote the splicing of its own mRNA (Fig. 3, *D* and *E*).

SPF30-mutant analysis showed that the expression of the ΔC5 deletion mutant (Δ218–238), which possessed RNA-binding properties as shown by EMSA analysis (Fig. 5*D*), did not cause SPF30 autoregulation, suggesting that the protein component required for the autoregulatory mechanism possibly binds to the aa 218–238 region of SPF30 and promotes splicing at the intron 3–exon 4a junction (Fig. 5*B*). Further identification of these factors and their roles in the autoregulatory mechanism of SPF30 will help elucidate key molecular mechanisms. In this study, the molecular mechanism of exon 4a inclusion via the SPF30 protein was revealed; however, it remains unclear whether direct binding of SPF30 to pre-mRNA is involved in the inclusion of cassette exons within intron 2. The results of previous surveys identified SPF30-binding RNA using enhanced cross-linking and immunoprecipitation (eCLIP), a UV-cross-linking-based method (35). However, because SPF30 transcripts with cassette exon and exon 4a are likely to be immediately degraded by NMD, SPF30 binding sites in SPF30 transcripts have not yet been detected. To identify transcripts that are degraded by NMD, additional UV-CLIP-based assays should be performed while inhibiting mRNA degradation with an NMD inhibitor, such as CHX.

Based on the results shown in Fig. 5*D*, we explored an SPF30–RNA-binding model using *in silico* protein–RNA docking simulations. Further simulations with SPF30 mutants lacking the kink-like structure (ΔC1–ΔC3) revealed different binding patterns with transcripts containing exon 4a versus WT SPF30 (Fig. S7, A–C). In particular, ΔC3 (containing a complete C-terminal α-helix of SPF30) was predicted to bind RNA through the α-helix of SPF30 (Fig. S7*C*). EMSA analysis of ΔC3 showed smeared band-shifts of BT-RNA, suggesting that the α-helix of SPF30 might have RNA-binding properties, although the interaction domain of SPF30 was insufficient for the binding observed between WT SPF30 and BT-RNA (Fig. 5*D*). In addition, a docking simulation using ΔC4 (Fig. S7*D*), containing half of the kink-like structure, predicted a specific binding mode similar to WT SPF30 and ΔC5, but EMSA analysis with ΔC4 showed only smear band-shifts of BT-RNA (Figs. 5*D*). This discrepancy may have arisen due to different approaches used to evaluate the contribution of RNA binding within the kink-like structure. Although the docking simulations suggested that the first half of the kink-like structure was sufficient for RNA binding, the entire kink-like structure appeared to be required for a particular binding mode, as indicated by the EMSA results. Integrating these results, we propose a reasonable model in which SPF30 binding to transcripts containing exon 4a requires both the C-terminal α-helix and the kink-like structure of SPF30.

## Experimental procedures

### Antibodies and reagents

The antibodies used in this study are listed in Supplementary Table S1. All reagents were purchased from Wako Pure Chemical Industries (Osaka, Japan) or Nacalai Tesque (Kyoto, Japan).

### Cell culture

Flp-In T-REx 293 (293FTR; Thermo Fisher Scientific) and HCT116 cells were cultured in Dulbecco’s modified Eagle’s medium with high glucose (Wako Pure Chemical Industries, Osaka, Japan). All cell lines were cultured at 37 °C under 5% CO_2_ in air. All media were supplemented with 10% fetal bovine serum (Nichirei Bioscience, Japan), streptomycin (0.1 mg/mL), and penicillin G (100 U/mL). Cycloheximide (Sigma-Aldrich, 239763-M) was used at a concentration of 50 ng/mL for the indicated periods. 5-Phenyl-indole-3-acetic acid (5-Ph-IAA; MedChemExpress, HY-134653) was used at concentrations of 0.25 μM for the indicated periods.

### Construction of expression plasmids

To construct a plasmid expressing SPF30-FLAG-6xHis (pcDNA5/FRT/TO SPF30(WT)-FH), DNA fragments encoding full-length SPF30 (NM_005871:174-887) were amplified using KOD Plus Neo (TOYOBO, Japan) with a primer set (KI-16/KI-17), using a modified pBICEP-CMV2 SPF30-FLAG as template DNA (26). The amplified DNA was inserted into the BamHI sites of pcDNA5/FRT/TO NVL2-FH (36) using the NEBuilder HiFi DNA Assembly Master Mix (New England Biolabs). Construction of the deletion mutant series of pcDNA5/FRT/TO SPF30-FH (ΔN, ΔTudor, ΔC3, ΔC4, and ΔC5) was performed using pcDNA5/FRT/TO SPF30(WT)-FH as template DNA as follows. For ΔN and ΔC1, DNA fragments were amplified using KOD FX Neo with primer sets (KI-101/KI-17 for ΔN and KI-16/KI-99 for ΔC1) and inserted within the BamHI sites of pcDNA5/FRT/TO NVL2-FH using the NEBuilder HiFi DNA Assembly Master Mix. For ΔTudor, ΔC4, and ΔC5, DNA fragments were amplified using KOD FX Neo with primer sets (KI-102/KI-103 for ΔTudor, KI-316/KI-319 for ΔC4, and KI-318/KI-319 for ΔC5), digested with DpnI and BamHI, and self-ligated with Ligation high Ver.2 (TOYOBO, Japan).

NanoLuc luciferase (Nluc)-expressing plasmid (pNlucP) was used for in vivo splicing assays. DNA fragments encoding Nluc were amplified using KOD FX Neo with a primer set (KI-231/KI-232) and a DNA template (pEX-A2J1-HA-NlucP, artificial DNA synthesis service in Eurofins Genomics). The amplified DNA was digested with BamHI and NotI and inserted into the BamHI and NotI sites of pEGFP-N1 (Clontech). For the construction of pNlucP Ex3-Int2-Ex3 WT, DNA fragments encoding SPF30(Ex3-Int3-Ex4) were amplified using KOD FX Neo with a primer set (KI-205/KI-226) and HCT116 genomic DNA as a DNA template, digested with HindIII and BamHI, and inserted into the HindIII and BamHI sites of pNlucP using Ligation high Ver.2. A mutant series of pNlucP Ex3-Int2-Ex3 (mut1, mut2, and mut3) was generated using the QuickChange protocol. DNA fragments were amplified using KOD FX Neo with primer sets (KI-305/KI-306 for mut1, KI-255/ KI-256 for mut2, and KI-257/ KI-258 for mut3) using pNlucP Ex3-Int2-Ex3 WT as a DNA template and digested with DpnI to degrade template DNA. To construct pNlucP HBB (Ex1-Int1-Ex2), DNA fragments were amplified using KOD FX Neo with a primer set (KI-251/ KI-252) using 293FTR genomic DNA as a DNA template, digested with HindIII and BamHI, and inserted into the HindIII and BamHI sites of pNlucP using Ligation high Ver.2.

Plasmids expressing SPF30 and its mutants in *Escherichia coli* were constructed as follows. DNA fragments were amplified using KOD FX Neo with primer sets (KI-143/KI-125) using a modified pBICEP-CMV2 SPF30-FLAG as template DNA (26) as a DNA template. The template DNA was digested with SacI and XbaI and inserted into the SacI and XbaI sites of pCold I (Clontech) with Ligation high Ver.2, which are referred to as pCold I SPF30 WT. To construct the deletion mutants of pCold I SPF30 (ΔC1, ΔC2, ΔC3, ΔC4, and ΔC5), DNA fragments were amplified using KOD FX Neo with primer sets (KI-143/KI-153 for ΔC1, KI-143/KI-219 for ΔC2, KI-143/KI-220 for ΔC3, KI-143/KI-316 for ΔC4, and KI-318/KI-320 for ΔC5) and pCold I SPF30(WT) as a DNA template. The amplified DNA was digested with SacI and BamHI and inserted into the SacI and BamHI sites of the pCold I SPF30 WT. To construct pCold I SPF30 ΔC6, DNA fragments were amplified using KOD FX Neo with primer sets (KI-370/KI-371), and pCold I SPF30 ΔC5 as DNA template, digested with DpnI and BamHI, and ligated with Ligation high Ver.2. All plasmid sequences were confirmed using DNA sequencing. Primers used in this study are listed in Supplementary Table S2. All plasmids used in this study are listed in Supplementary Table S3.

### Construction of doxycycline-inducible cell lines

293FTR cells expressing the target protein with doxycycline treatment were established using the Flp-In T-REx Expression system (Thermo Fisher Scientific, USA), as described previously (17). Briefly, 293FTR cells were transfected with pcDNA5/FRT/TO SPF30-FLAG or its mutants and pOG44 (Thermo Fisher Scientific, USA) using Lipofectamine 2000 reagent (Invitrogen, USA) according to the manufacturer’s instructions. Clonal cells were obtained by selection in culture medium containing 100 μg/mL hygromycin B (Wako Pure Chemical Industries, Osaka, Japan). The cells were treated with 1 μg/mL doxycycline for the indicated times to express SPF30-FLAG or its mutants.

### Western blotting

Western blotting was performed as previously described (37). Chemiluminescent signals of the protein bands were detected using FUSION Solo (Vilber Lourmat, Germany). The antibodies used in this study are listed in Supplementary Table S1.

### Immunocytostaining

Cells cultured on collagen-coated culture slides were washed with phosphate-buffered saline (PBS) and fixed with 4% paraformaldehyde in PBS for 10 min at 25 °C. After washing twice with PBS containing 0.05% (v/v) Tween 20 (PBST), the cells were permeabilized with PBS containing 0.1% (v/v) Triton X-100 (SIGMA) for 5 min at 25 °C and then washed again with PBST. The cells were incubated with blocking buffer (3% (w/v) BSA/PBS) for 1 h and then incubated with an appropriate primary antibody in 1% (w/v) BSA/PBS overnight at 4 °C. After three 10-min washes with PBST, the cells were incubated with a fluorescence-conjugated secondary antibody for 1 h at 25 °C. Finally, after three 10-min washes with PBST, the cells were mounted using a mounting medium with Hoechst. Fluorescence imaging was performed using a BZ-700 All-in-one microscope (Keyence, Osaka, Japan) with a 100× objective. Antibodies used in this study are listed in Supplementary Table S1.

### RT-qPCR and RT-PCR analysis

Total RNA was isolated using TRIzol Reagent (Thermo Fisher Scientific) or Sepazol-RNA I Super G (Nacalai Tesque, Kyoto, Japan), according to the manufacturer’s instructions. For RT-qPCR analysis, reverse transcription was performed using the PrimeScript RT reagent kit (Perfect Real Time; Takara Bio, Japan) with oligo dT and random hexamer. Quantitative PCR was performed using the THUNDERBIRD Next SYBR qPCR Mix (TOYOBO, Japan) with the indicated primer sets on a CFX Connect Real-Time PCR system (Bio-Rad, CA, USA). For RT-PCR analysis, reverse transcription was performed using ReverTra Ace (TOYOBO, Japan) with oligo dT primers. RT-PCR was performed using KOD FX Neo (TOYOBO, Japan) with the indicated primer sets on a GeneAmp PCR System 9700 thermocycler (Applied Biosystems, USA). For RT-qPCR, primer sets of KI-39/KI-40 for SPF30 3’UTR, KI-41/KI-42 for SPF30 CDS, KI-78/KI-79 for SPF30 Int2/Ex3, KI-97/KI-130 for SPF30 Ex3/Ex4a, and KI-173/KI-174 for ACTB were used. For RT-PCR, primer sets of KI-43/KI-44 for SPF30 Ex1/Ex6, KI-43/KI-42 for SPF30 Ex1/Ex4, KI-41/KI-44 for SPF30 Ex4/Ex6, KI-43/KI-97 for SPF30 Ex1/Ex4a, and KI-58/KI-59 for ACTB were used. Primers used in this study are listed in Supplementary Table S2.

### Library preparation for deep sequencing and analysis

293FTR cells expressing SPF30-FLAG were treated with or without 1 µg/ml doxycycline for 48 h and treated with 50 µg/ml cycloheximide for 4 h before harvesting. Cells were harvested and total RNA was extracted as described above. cDNA was prepared using Superscript III (Invitrogen) according to the manufacturer’s instructions. The exon 1 to exon 4 region (111-690) of SPF30 was amplified with primer sets KI-178/KI-179 using KOD FX Neo, and the amplified DNA was separated by 1.5% agarose gel electrophoresis. PCR products with 400 bp (without the cassette exon) or 600 bp (with the cassette exon) in lanes 2 and 4 in Fig. S2, respectively, were excised and extracted using the MinElute PCR Purification kit (Qiagen). DNA libraries derived from each DNA fragment (25 ng) for deep sequencing were prepared using a KAPA Hyper Prep kit for Illumina (Kapa Biosystems). Briefly, after end repair and A-tailing, the KAPA Universal Adapter (Kapa Biosystems) was ligated to the PCR products using DNA ligase. Next, ligated DNAs were amplified using KAPA UDI Primer Mixes (Kapa Biosystems) and size selected with KAPA HyperPure Beads (Kapa Biosystems). The quantity and quality of the PCR products or libraries were determined using a Qubit 4.0, Qubit dsDNA HS Assay kit, and Bioanalyzer. Sequencing of the prepared libraries by NovaSeq X Plus (Illumina) was performed using the commercial services (Gigabase Reading service, PE-150 or PE-250) of Nippon Gene (Tokyo, Japan). The libraries from 400 bp and 600 bp PCR products were sequenced with paired-end 150 bp reads and paired-end 250 bp reads, respectively.

Adaptor sequences and low-quality bases were trimmed from raw reads using the Fastp trimmer, and quality checks were subsequently conducted with FastQC. For analysis of the 400 bp PCR product, reads including the sequences of Ex4 (GTGTTATGAAGC), Ex3/Ex4 (TGGACAGTGTT), and Ex3/Ex4a (TGGACACCATGATA) were counted using the Seqkit tool with the command “seqkit-grep” (http://bioinf.shenwei.me/seqkit)(***38). For the analysis of the PCR product with 600 bp, reads including the sequences of cassette exon (cEx) and Ex4 (TAATGGAAGT) were extracted as the raw reads, and subsequently, the reads including the sequences of Ex4 (GTGTTATGAAGC), Ex3/Ex4 (TGGACAGTGTT), and Ex3/Ex4a (TGGACACCATGATA) were counted by Seqkit tool with the command “seqkit-grep.” The ratio of read counts for Ex3/Ex4 or Ex3/Ex4a per Ex4 was calculated.

### Construction of SPF30 knockdown-inducible cell line

For producing the Cas9-expressing vector containing the sequence of sgRNA targeting human SPF30, we referred to the method described previously (39). Briefly, oligo dsDNA, which is composed of sense-oligo and antisense-oligo ssDNAs (SM-1/SM-2), was inserted into peSpCas9(1.1)-2×sgRNA (Addgene#80768) in the solution containing BpiI (BbsI; Thermo Scientific) and Ligation high Ver.2 (Toyobo, Japan). The procedure for generating donor plasmids for the addition of AID tags to endogenous genes referred to the method described previously (40). Briefly, genomic PCR was first performed using genomic DNA extracted from HCT116 cells as a template and primers with SacI and KpnI sites added to the 5’ end of each (SM-3/SM-4) for amplifying the homology arm (HA) sequence against the SPF30 gene. This PCR amplicon was then digested with SacI and KpnI and inserted into the SacI/KpnI sites of the pBluescript II SK(+) vector. Next, the HA-inserted vector was subjected to inverse-PCR using primer set (SM-5/SM-6) for introducing the BamHI site just before the terminal codon of the SMNDC1 gene as the first step, and then using primer set (SM-7/SM-8) for mutating to the CRISPR-Cas9 targeting sequence as the second step. After two rounds of inverse-PCR, the intermediate plasmid was digested using BamHI and inserted into the BamHI site of pMK293 (RIKEN DNA BANK) or pMK393 (RIKEN DNA BANK), which contain mAID-mCherry tag and antibiotics resistance markers. In plasmid construction, PCR reactions were performed using KOD FX Neo DNA polymerase (Toyobo, Japan). All primers were listed in Supplementary Table S2.

For mAID-mCherry-tagging of the endogenous protein using the CRISPR/Cas9 system, HCT116/OsTIR1 F74G parental cells (30) were treated with 40 µM SCR7. Four hours later, 1 µg of guide RNA and Cas9 all-in-one plasmid and two homology-dependent repair (HDR) donor plasmids (a total of 2 μg) were transfected into a 60-mm dish, and further 24h later, cells were transferred to a 100-mm dish containing 100 µg/mL hygromycin B and 15 µg/mL blasticidin S. After approximately 14 days of selection, a single cell with mCherry fluorescence was sorted to 96-well plates using a MoFlo XDP cell sorter (Beckman Coulter). Screening of alleles and expressed proteins in cloned cells was performed by genomic PCR and western blotting, respectively. For inducing degradation of an endogenous protein fused with mAID-mCherry tag, 5-Ph-IAA dissolved in DMSO was added directly to the culture medium at the indicated concentration.

### Luciferase assay

pCMV-Ex3-Int3-Ex4-NLucP or its mutants (20 ng) and pGL4.54 [luc2/TK] as an internal control (80 ng; Promega, USA) were transfected into SPF30-mAID cells on a 12-well plate using ViaFect (Promega, USA) according to the manufacturer’s instructions. pCMV-HBB-NLucP was used as a control for pCMV-Ex3-Int3-Ex4-NLucP mutants. After 24 h of transfection, cells were treated with or without 5-Ph-IAA (0.25 μM). After 24 h of transfection, cells were harvested and lysed with 300 μl of 1× Passive Lysis Buffer (Promega, USA), and then 100 μl of each lysate was subjected to a luciferase assay. Luciferase assays were performed using the Dual Glo Luciferase Assay System (Promega, USA) and the GloMax Discover System (Promega, USA) according to the manufacturer’s instructions.

### in vitro protein–RNA-binding assay

SPF30-FLAG or its mutant-inducible cells were treated with doxycycline for 48 h and harvested with ice-cold PBS. Cells were lysed with lysis buffer (50 mM Tris–HCl, pH 7.4, 150 mM NaCl, 0.5% IGEPAL CA-630) containing 1 mM PMSF, 1 μg/ml aprotinin, 1 μg/ml pepstatin, and 10 μg/ml leupeptin for 10 min on ice. After centrifugation at 20,000 × g for 10 min at 4 °C, the supernatant was collected and used as cell extract. The cell extract was incubated with 25 µl FLAG magnetic beads and 1 pmol BT-RNA biotinylated RNA (WT or mut1) for 4-h rotation at 4 °C. The beads were washed five times with lysis buffer and then exposed to UV light at 400 mJ/cm^2^ using the CX-2000 UV crosslinker (UVP) for cross-linking of the binding of SPF30-FLAG and BT-RNA. The beads were resuspended in 1 ml high-salt RIPA buffer (50 mM Tris–HCl, pH 7.4, 400 mM NaCl, 1% IGEPAL CA-630, 0.1% SDS, 0.5% sodium deoxycholate) containing 1 mM PMSF and rotated for 1 h at 4 °C. After washing five times with 1 ml of high-salt RIPA buffer, SPF30-FLAG and BT-RNA were eluted with SDS-PAGE sample buffer after incubation for 3 min at 65 °C. SPF30-FLAG and BT-RNA were separated by SDS-PAGE on a 7% acrylamide gel (19:1)/Bis-Tris buffer in MOPS running buffer and electrophoretically transferred to a nitrocellulose membrane (Protran BA 85, 0.45 µm, Whatman). The membrane was then exposed to UV light (120 mJ/cm^2^). BT-RNA was detected using streptavidin-horseradish peroxidase (HRP; Thermo Fisher Scientific), and SPF30-FLAG was detected using an anti-SPF30 antibody or anti-FLAG (M2) antibody. Chemiluminescence signals were visualized using FUSION Solo. The BT-RNA sequence used in this study is described in Fig. 4A and Supplementary Table S2.

### RNA immunoprecipitation (RIP) assay

293FTR cells expressing SPF30 or not were cultured in a 150-mm Petri dish and harvested with ice-cold PBS. Cells were washed twice with ice-cold PBS and lysed with lysis buffer [50 mM Tris–HCl, pH 7.4, 150 mM NaCl, 0.5% (v/v) IGEPAL CA-630 (SIGMA)] containing 1mM PMSF, 1 μg/ml aprotinin, 1 μg/ml pepstatin, and 10 μg/ml leupeptin for 10 min at 4 °C. After centrifugation at 20,000 × g for 10 min at 4 °C, the supernatant was rotated with 20 μl of anti-FLAG M2 agarose beads (Sigma-Aldrich) for 2 h at 4 °C. The agarose beads were washed five times with lysis buffer and eluted with 150 μl of protein–RNA extraction buffer (7 M urea, 350 mM NaCl, 1% SDS, 10 mM Tris–HCl, pH 8.0, 10 mM EDTA, and 2% 2-mercaptoethanol) for 5 min at 25 °C. Protein and RNA components were extracted as previously described (37) and subjected to western blotting or RT-PCR, using the indicated primer sets.

### Purification of recombinant proteins

6xHis-tagged SPF30 (6xHis-SPF30) or its mutants (WT, ΔC1, ΔC2, ΔC3, ΔC4, ΔC5, or ΔC6) were expressed in *Escherichia coli* BL21-CodonPlus(DE3)-RIPL (Stratagene) in 200 ml of LB medium using pCold I DNA (Takara Bio, Japan) via 15 h induction at 16 °C in the presence of 0.1 mM Isopropyl β-D-1-thiogalactopyranoside. The collected bacteria expressing 6xHis-SPF30 (WT or its mutants) were resuspended in 10 ml of lysis buffer (10 mM Tris–HCl pH 8.0, 150 mM NaCl, 1% Triton X-100) containing 1 mM DTT and 1 mM PMSF, and lysed by sonication using a Bioruptor 250 (highest setting, six 30-s pulses with 30-s intervals, 4 °C; CosmoBio, Japan). After centrifugation at 20,000 × *g* for 10 min at 4 °C, the supernatant was added to 200 μl of Ni-NTA resin (Qiagen) and rotated at 4 °C for 1 h. The resin was washed five times with lysis buffer, and the recombinant proteins were eluted with elution buffer (10 mM Tris–HCl pH 8.0, 150 mM NaCl, 250 mM imidazole). 6xHis-SPF30 or its mutants were purified by size-exclusion chromatography using Superose 6 (GE Healthcare) in an AKTA Explorer system (GE Healthcare). The protein concentrations of the purified proteins were measured using Pierce 660 nm Protein Assay Reagent (Thermo Fisher Scientific) and bovine serum albumin.

### Electrophoretic mobility shift assay (EMSA)

Biotin-labeled synthetic RNA (BT-RNA) was purchased from Fasmac Co. Ltd. (Japan). The indicated amounts of recombinant protein and 20 fmol of BT-RNA were incubated in 12 μl binding buffer (40 mM Tris–HCl, pH 7.4, 30 mM KCl, 1 mM MgCl_2_, 0.01% IGEPAL CA-630 (octylphenoxy poly(ethyleneoxy)ethanol) for 30 min at 25 °C. RNA–protein complexes were separated by 7% non-denaturing polyacrylamide gel electrophoresis at 100 V for 90 min in 0.5× Tris-borate/EDTA buffer (44.5 mM Tris-borate and 1 mM EDTA) and transferred to a Hybond N+ membrane (Cytiva, USA). The membrane was sequentially dried and UV-crosslinked using a CX-2000 UV crosslinker (UVP) at 120 mJ/cm^2^. BT-RNA was detected using a Chemiluminescent Nucleic Acid Detection Module Kit (Thermo Fisher Scientific), according to the manufacturer’s instructions. Signals of BT-RNA were visualized by FUSION Solo. The BT-RNA sequence used in this study is described in Fig. 4A and Supplementary Table S2.

### in Silico Protein–RNA docking analysis

All docking calculations were performed using the HDOCK web server, which is designed for protein– RNA docking (32, 33). The HDOCK server relies on a hybrid algorithm that combines template-based modeling with *ab initio*-free docking. The structure of SPF30 was predicted using AlphaFold2 (Fig. S4A)(31), while the dsRNA structure was modeled using the RNAfold web server (http://rna.tbi.univie.ac.at/cgi-bin/RNAWebSuite/RNAfold.cgi) (34) and RNA tertiary structure was obtained using the RNAcomposer web server (https://rnacomposer.cs.put.poznan.pl) (41, 42). All docking results discussed in this work are based on the top-ranked (rank 01) predicted structures. Additionally, all mutant structures of SPF30 (ΔC1, ΔC2, ΔC3, ΔC4, and ΔC5) were generated by fixing the backbone of the SPF30 WT and introducing mutations solely to the side chains. The structure prediction using AlphaFold2 was performed with ColabFold v1.5.5. The tertiary structures of protein and RNA were made with VMD software (43).

## Data availability

All data for this publication are included in the article or are available from the corresponding authors upon reasonable request.

## Statistical analysis

Statistical analyses were performed with GraphPad Prism 10.4.0. Comparisons between indicated groups were performed using Welch’s test, Tukey’s multiple comparisons test, or Dunnett’s test. P values < 0.05 were considered statistically significant. *P < 0.05, **P < 0.01, ***P < 0.001, ****P < 0.0001, and ns for P ≥ 0.05.

## Supporting information

This article contains the supporting information.

## Author contributions

**Keiichi Izumikawa:** Writing – original draft, Investigation, Validation, Methodology, Investigation, Conceptualization, Project administration, Supervision, Funding acquisition. **Tatsuya Shida:** Writing – original draft, Investigation, Conceptualization, Resources. **Yuuka Onodera:** Investigation, Resources. **Yuito Tashima:** Investigation, Resources. **Sotaro Miyao:** Resources. **Tomomi Suda:** Methodology. **Yasuyuki Suda:** Methodology. **Minoru Sugihara:** Writing – original draft, Methodology, Investigation, Funding acquisition. **Tamotsu Noguchi:** Methodology. **Masami Nagahama:** Writing – original draft, Conceptualization, Supervision, Funding acquisition.

## Acknowledgments

We thank Dr. Yohei Kato and Prof. Kazuhisa Nakayama (Kyoto University, Japan) for the initial advice regarding genome editing experiments. The single cell sorting was supported by Stem Cell Laboratory, Advanced Technology Laboratory, Medical Research Laboratory, Institute for Integrated Research, Institute of Science Tokyo. We would like to thank Editage (http://www.editage.jp) for English language editing.

## Funding and additional information

This work was partly supported by Grants-in-Aid for Scientific Research (C) [grant numbers 19570180 and 24590101 to M.N.] from the Japan Society for the Promotion of Science (JSPS). This work was supported by Joint Research Program in Meiji Pharmaceutical University (JRP24MPU to K.I, M.S.).

## Abbreviations

aa: amino acid
AID: auxin-inducible degron
ALS: amyotrophic lateral sclerosis
BT-RNA: biotinylated RNA
CDS: coding sequence
CHX: cycloheximide
Dox: doxycycline
eCLIP: enhanced cross-linking and immunoprecipitation
EMSA: electrophoretic mobility shift assay, ex: exon
HHB: human hemoglobin subunit beta
HRP: horseradish peroxidase
IAA: 5-phenyl-1H-indole-3-acetic acid
mCherry: monomeric Cherry
Nluc: NanoLuc Luciferase
NMD: nonsense-mediated mRNA decay
nt: nucleotide
OsTIR1(F74G): F74G variant of the Arabidopsis thaliana TIR1 protein
PC: Parental HCT116 cell
PTC: premature termination codon
RT-qPCR: quantitative RT-PCR
SC35: serine/arginine-rich splicing factor 2
SD: standard deviation
snRNP: small nuclear ribonucleoprotein
SPF30-FLAG: C-terminally FLAG-tagged and 6xHis-tagged variant of SPF30
SPF30-mAC: SPF30 variant containing AID and mCherry tags
SPF30-mAID: clonal cell line expressing an SPF30 containing AID and mCherry tags
SPF30: motor neuron-related-splicing factor 30
TDP-43: TAR DNA-binding protein 43
UTR: untranslated region
WT: wild-type

