## Supporting information for "Molecular basis of the autoregulatory mechanism of motor neuron-related splicing factor 30"

### Supporting Tables

#### Supplementary Table S1. List of antibodies used in this study.

| Antibody | Host/isotype | Supplier | Product code | Used for |
|---|---|---|---|---|
| β-actin (AC-15) | Mouse / IgG1 | Santa Cruz Biotechnology | sc-69879 | WB |
| GAPDH (6C5) | Mouse / IgG1 | Ambion | AM4300 | WB |
| FLAG (M2) | Mouse / IgG1 | SIGMA | F3165 | WB |
| FLAG | Rabbit | SIGMA | F7425 | ICS |
| SPF30 (SMNDC1) | Rabbit | INVITROGEN | PA5-31148 | WB, ICS |
| SC-35 | Mouse / IgG1 | SIGMA | SAB4200725 | ICS |
| Anti-mouse IgG, HRP-linked Antibody | Horse | Cell Signaling Technology | #7076 | WB |
| Anti-Rabbit IgG (HTL)-HRP Conjugate | Goat | Bio-Rad | #1706515 | WB |
| Anti-mouse IgG (HTL)-HRP Conjugate | Goat | Bio-Rad | #1706515 | WB |
| Anti-Goat IgG –Peroxidase antibody | Rabbit | SIGMA | A5420 | WB |
| Anti-mouse IgG (Fc BP)-HRP Conjugate | | Santa Cruz Biotechnology | sc-525409 | WB |
| Goat anti-Mouse IgG (H+L) Cross-Adsorbed Secondary Antibody, Alexa Fluor 488 | Goat | Themo Fisher Scientific | A-11001 | ICS |
| Goat anti-Rabbit IgG (H+L) Cross-Adsorbed Secondary Antibody, Alexa Fluor 594 | Goat | Themo Fisher Scientific | A-11012 | ICS |

### Supplementary Table S2. List of primers used in this study.

| Primer Name | Sequence 5' --> 3' | Used for |
|---|---|---|
| KI-173 | CATGTACGTTGCTATCCAGGC | ACTB-Fw, qPCR |
| KI-174 | CTCCTTAATGTCACGCACGAT | ACTB-Rv, qPCR |
| KI-58 | AACAAGATGAGATTGGCA | ACTB-Fw, RT-PCR |
| KI-59 | GACCAAAAGCCTTCATACAT | ACTB-Rv, RT-PCR |
| KI-39 | CTAGCAATTCATTGGAACACCA | SPF30 3'UTR(Exon 6)-Fw, qPCR |
| KI-40 | GCCTGGGAAACAAGTTATTGAG | SPF30 3'UTR(Exon 6)-Rv, qPCR |
| KI-41 | AGCGGAGATTGAGGAGATAGATG | SPF30 CDS(Ex4)-Fw, qPCR, RT-PCR |
| KI-42 | GGTTCAACAGTGGAGTCACTTC | SPF30 CDS(Ex4)-Rv, qPCR, RT-PCR |
| KI-43 | TTTCCTGCTACTGCTACTGCTG | SPF30 Ex1-Fw, RT-PCR |
| KI-44 | CTGCAGAGATGAAATCCAACAG | SPF30 Ex6-Rv, RT-PCR |
| KI-78 | TTCCTCCCTCAAGTCTGAGTTC | SPF30 Int2-Rv, qPCR |
| KI-79 | TACAAAGCTCAGCTCCAGCA | SPF30 Ex2-Fw, qPCR |
| KI-97 | ACACCTGTAAAGATATCATGGTGTC | SPF30 Ex4a-Rv, qPCR, RT-PCR |
| KI-130 | GAACTAACCAAAGACCTTCTGTCAA | SPF30 Ex3-Fw, qPCR |
| KI-16 | AGCTTGGTACCGAGCTCGACCGCCATGTCAGAGGATTTAGCAAAGCAG | Construction of pcDNA5FRT/TO SPF30(WT, ΔC1)-FLAG-6His |
| KI-17 | TCCTTGTAATCGGTACCGGCTTGAGGCATCAAATGCCTGACATT | Construction of pcDNA5FRT/TO SPF30(WT, ΔN)-FLAG-6His |
| KI-101 | AGCTTGGTACCGAGCTCGGATCCACCGCCATGTCTACTCAACCTACTCATTCATGG | Construction of pcDNA5FRT/TO SPF30(ΔN)-FLAG-6His |
| KI-99 | TCCTTGTAATCGGTACCGGCTATTCTCTGAGCTTTTTTCAAAGC | Construction of pcDNA5FRT/TO SPF30(ΔC1)-FLAG-6His |
| KI-102 | TATAGGATCCAGAAGCAAAACTGTCTGAACT | Construction of pcDNA5FRT/TO SPF30(ΔTudor)-FLAG-6His |
| KI-103 | TATAGGATCCAAGGAGGACAGTGGCAACAAA | Construction of pcDNA5FRT/TO SPF30(ΔTudor)-FLAG-6His |
| KI-316 | TATAggatccCACACTCTCAGGTGAAGCAAAAATA | Construction of pcDNA5/FRT/TO SPF30(ΔC4)-FLAG-6His,<br>Construction of pCold-I SPF30(ΔC4) |
| KI-319 | tataggatccGCCGGTACCGATTACAAGGACGAC | Construction of pcDNA5/FRT/TO SPF30(ΔC4, ΔC5)-FLAG-6His |
| KI-318 | TATAggatccAGCAATTCCACAGGTTCCTACTCC | Construction of pcDNA5/FRT/TO SPF30(ΔC5)-FLAG-6His,<br>Construction of pCold-I SPF30(ΔC5) |
| KI-231 | tataGGATCCAtacccttatgacgtcccggactac | Construction of pNanoLucP SPF30(Ex3-Int3-Ex4)(WT) |
| KI-232 | TATAgcggccgcTTATGACGTTGATGCGAGCTGAAG | Construction of pNanoLucP SPF30(Ex3-Int3-Ex4)(WT) |
| KI-205 | TATAaagcttACCGCCATGGAACTAACCAAAGACCTTCTGTCA | Construction of pEGFP-N1 SPF30(Ex3-Int3-Ex4) |
| KI-226 | TATAggatccggGCCATAACCAGCAAAGGTGATTG | Construction of pEGFP-N1 SPF30(Ex3-Int3-Ex4) |
| KI-305 | TGCTGGTGGTCGAGATTGAGGAGATAGATGAAGAA | Construction of pNanoLucP SPF30(Ex3-Int3-mEx4)(mut1) |
| KI-306 | CGACCACCAGCAACCTGTAAAGATATCATGGCTGT | Construction of pNanoLucP SPF30(Ex3-Int3-mEx4)(mut1) |
| KI-255 | TTCCCACCCTCAGGTGTTATGAAGCGGAGATTGAG | Construction of pNanoLucP SPF30(Ex3-Int3-mEx4a)(mut2) |
| KI-256 | AGGGTGGGAAATGGCTGTAAGCAGGTGAGTAAAGG | Construction of pNanoLucP SPF30(Ex3-Int3-mEx4a)(mut2) |
| KI-257 | TTCCCACCCTCAGCCATGATATCTTTACAGGTGTT | Construction of pNanoLucP SPF30(Ex3-mInt3-Ex4)(mut3) |
| KI-258 | AGGGTGGGAAAGTAAAGGTTTAGTACTCAGCCTAC | Construction of pNanoLucP SPF30(Ex3-mInt3-Ex4)(mut3) |
| KI-251 | tataAAGCTTACCATGGTGCATCTGACTCCTGAGG | Construction of pNanoLucP HBB(Ex1-Int1-Ex2) |
| KI-252 | tataGGATCCTTAGGGTTGCCCATAACAGCATCA | Construction of pNanoLucP HBB(Ex1-Int1-Ex2) |
| KI-143 | tataGAGCTCATGTCAGAGGATTTAGCAAAGCAG | Construction of pCold-I SPF30(WT, ΔC1, ΔC2, ΔC3, ΔC4) |
| KI-125 | TATATCTAGACTACTTGTCATCGTCGTCCTTGTA | Construction of pCold-I SPF30(WT) |
| KI-153 | TATAggatccTATTCTCTGAGCTTTTTTCAAAGC | Construction of pCold-I SPF30(ΔC1) |
| KI-219 | TATAggatccAGAATAGGCTCTGTTGTTGAATTG | Construction of pCold-I SPF30(ΔC2) |
| KI-220 | TATAggatccCCTCTTTACCTGGCCTTTTTTGTT | Construction of pCold-I SPF30(ΔC3) |
| KI-320 | GATATCAGTCGACGGATCCGGTACCGATTA | Construction of pCold-I SPF30(ΔC5) |
| KI-370 | CCTCTTTACCTGGCCTTTTTTGTT | Construction of pCold-I SPF30(ΔC6) |
| KI-371 | ACTGGTAAAGTTGGAGTAGGAACC | Construction of pCold-I SPF30(ΔC6) |
| KI-178 | CCACTACCACTGGGCTCATT | Library preparation for deep sequencing, SPF30-Exon1-Fw |
| KI-179 | CGGTGCCATTTTCTTCATCT | Library preparation for deep sequencing, SPF30-Exon4-Rv |
| SM-1 | CACCAAAGTTGGAGTAGGAACCTG | Construction of peSpCas9(1.1)-2×sgRNA SPF30 |
| SM-2 | AAACCAGGTTCCTACTCCAACTTT | Construction of peSpCas9(1.1)-2×sgRNA SPF30 |
| SM-3 | ACTGGAGCTCACAGGCCCCACAGTAAACCACA | Construction of pBlueScript II SK (+)-SPF30-HA |
| SM-4 | AGCTGGTACCGCAGGTTGGTGTTTTTGCCTTGG | Construction of pBlueScript II SK (+)-SPF30-HA |
| SM-5 | GGATCCTAAAGGGCTTTACATTTACC | Construction of pBlueScript II SK (+)-SPF30-HA |
| SM-6 | TTGAGGCATCAAATGCCTGACATTG | Construction of pBlueScript II SK (+)-SPF30-HA |
| SM-7 | AAGGTAGGTGTTGGAACCTGTGGAATTGCTG | Construction of pBlueScript II SK (+)-SPF30-HA |
| SM-8 | ACCAGTCACACTCTCAGGTGAAGC | Construction of pBlueScript II SK (+)-SPF30-HA |

**Supplementary Table S3. List of plasmids used in this study**.

| Plasmid | Methods for construction |
|---|---|
| pcDNA5/FRT/TO DAP-Ex2-Int2-Ex3 | Izumikawa et al. 2018 |
| pcDNA5FRT/TO FLAG-Chtop | Izumikawa et al. 2016 |
| pOG44 | Themo Fisher Scientific (USA) |
| Modified pBICEP-CMV2 SPF30-FLAG | Ishida et al. 2021 |
| pcDNA5FRT/TO SPF30(WT)-FLAG-6His | |
| pcDNA5FRT/TO SPF30(ΔN)-FLAG-6His | |
| pcDNA5FRT/TO SPF30(ΔTudor)-FLAG-6His | |
| pcDNA5FRT/TO SPF30(ΔC1)-FLAG-6His | |
| pcDNA5FRT/TO SPF30(ΔC4)-FLAG-6His | |
| pcDNA5FRT/TO SPF30(ΔC5)-FLAG-6His | |
| pCold-I SPF30(WT) | |
| pCold-I SPF30(ΔC1) | |
| pCold-I SPF30(ΔC2) | |
| pCold-I SPF30(ΔC3) | |
| pCold-I SPF30(ΔC4) | |
| pCold-I SPF30(ΔC5) | |
| pCold-I SPF30(ΔC6) | |
| pEGFP-N1 SPF30(Ex3-Int3-Ex4) | |
| pNanoLucP SPF30(Ex3-Int3-Ex4)(WT) | |
| pNanoLucP SPF30(Ex3-Int3-mEx4)(mut1) | |
| pNanoLucP SPF30(Ex3-Int3-mEx4a)(mut2) | |
| pNanoLucP SPF30(Ex3-mInt3-Ex4)(mut3) | |
| pNanoLucP HBB(Ex1-Int1-Ex2) | |
| peSpCas9(1.1)-2×sgRNA | Addgene#80768 |
| peSpCas9(1.1)-2×sgRNA SPF30 | |
| pBlueScript II SK (+) | - |
| pBlueScript II SK (+) SPF30-HA | |
| pMK293 | RIKEN DNA BANK |
| pMK393 | RIKEN DNA BANK |
| pBlueScript II SK (+) SPF30-HA-pMK293 | |
| pBlueScript II SK (+) SPF30-HA-pMK393 | |

#### Supporting Figures

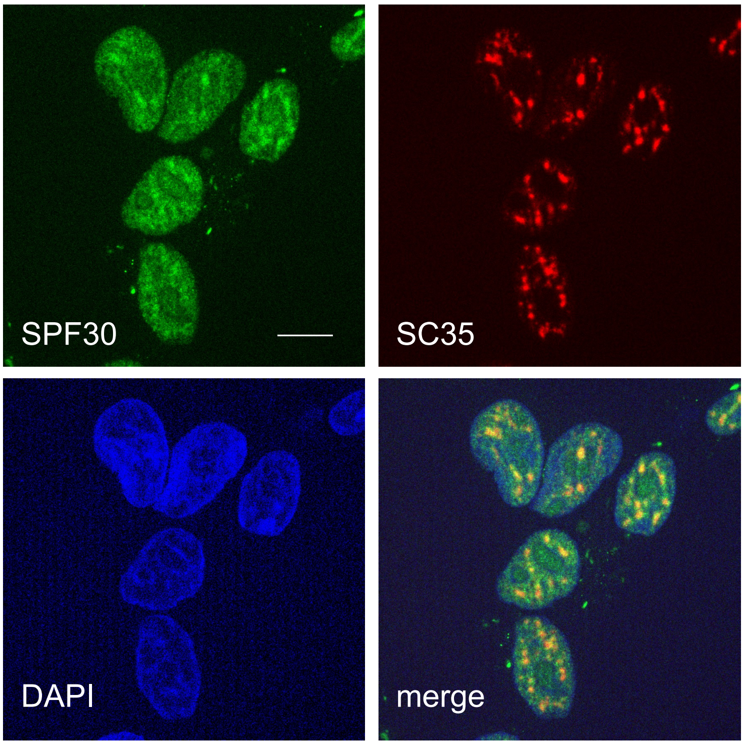


**Supplementary Figure S1.**

Immunofluorescent images of SPF30 (Green) and SC35 (Red) in 293FTR cells. In 293FTR cells, SPF30 and SC35 were stained with anti-SPF30, and anti-SC35 antibodies as the first antibodies, and stained with Alexa488-conjugated anti-rabbit IgG and Alexa595-conjugated anti-mouse IgG as the secondary antibodies. SC35 was used as nuclear speckles marker. DAPI was used as nuclear marker. Scale bar; 10 um.

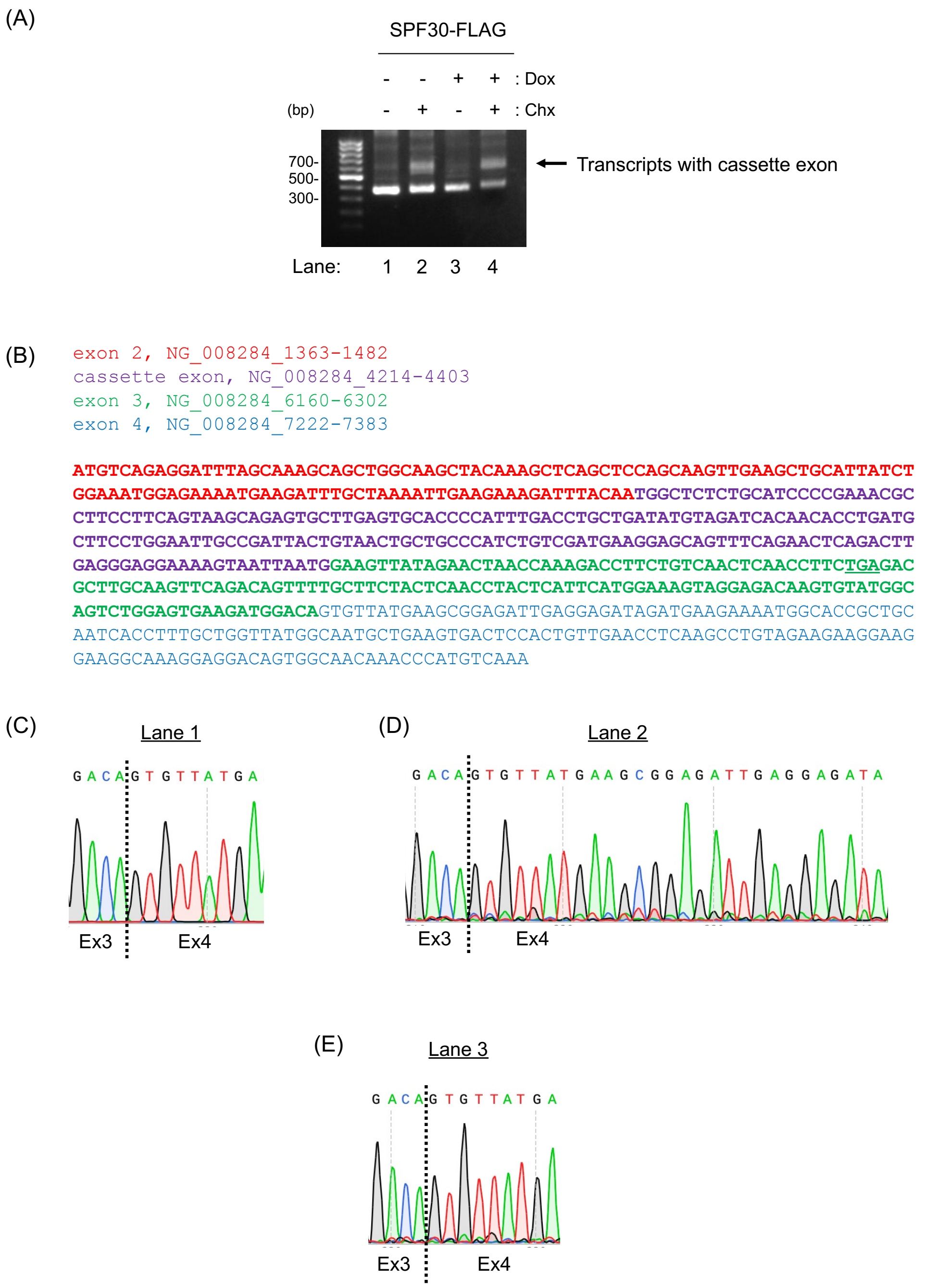


**Supplementary Figure S2.**

SPF30 transcripts with the cassette exon within intron 2 and exon 4a were generated upon CHX or SPF30 overexpression. (A) RT-PCR analysis of SPF30 transcripts for the deep sequencing. SPF30-FLAG inducible cells were treated with/without Dox for 48 h and CHX for 4 h, and total RNA was extracted. Endogenous SPF30 transcripts were detected by RT-PCR using exon 1 and exon 4 primer sets (KI-178/KI-179). (B) Sequence of SPF30 transcripts with the cassette exon, from exon2 to exon4, was annotated by the deep sequencing. (C-E) The PCR fragments with 500 bp, amplified by Ex1/Ex4, in lane2 (C), lane3 (D), and lane 4 (E) of Figure 2B were annotated by Sanger sequencing. Wave data at the junction of exon 3 and exon 4 from each Sanger sequencing were presented.

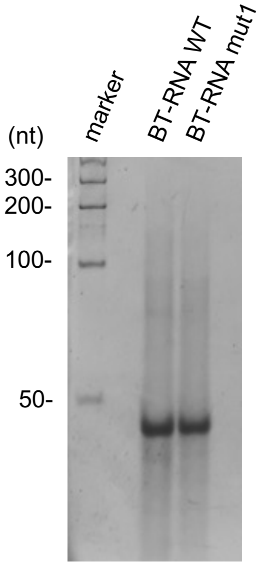


**Supplementary Figure S3.**

The biotinylated synthetic RNAs (BT-RNA WT and BT-RNA mut1) used in *in vitro* binding assays was separated by a denaturing urea-PAGE, and stained with SYBR gold. 1 pmol of RNAs was loaded to the denaturing urea-PAGE.

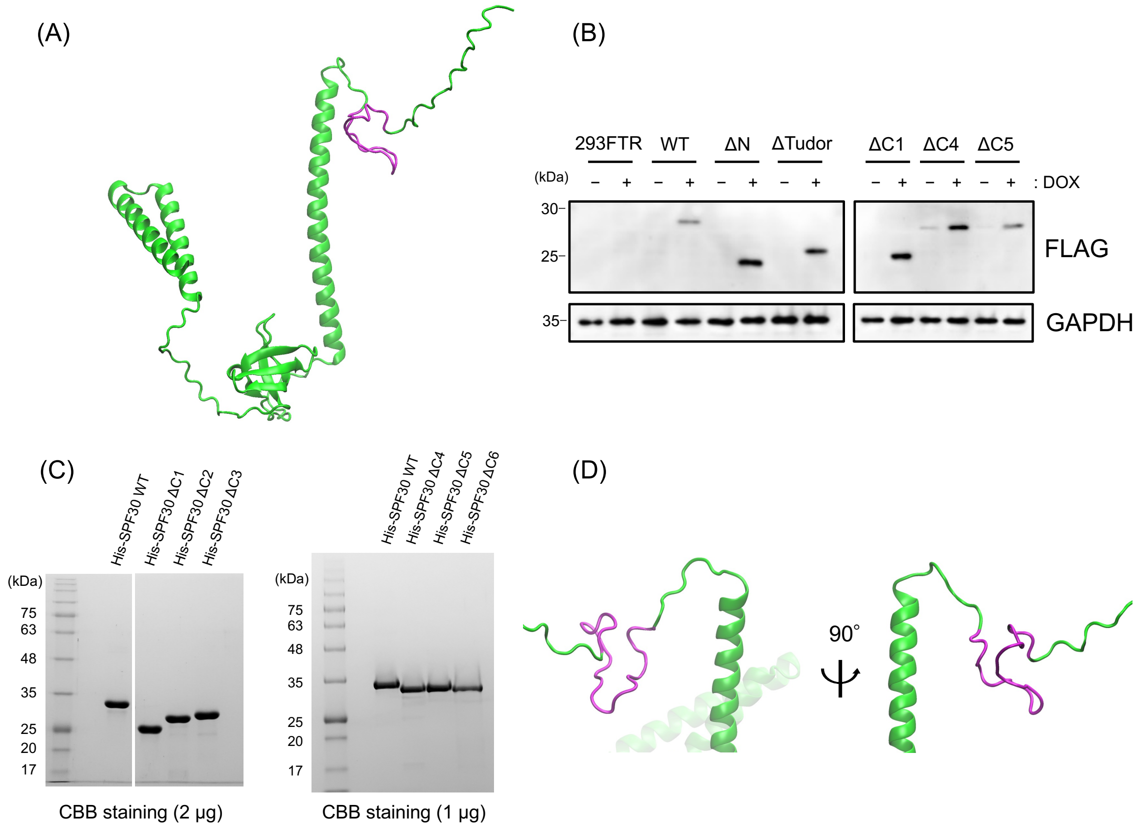


**Supplementary Figure S4.**

A structure model of SPF30 by Alphafold2. (A) A structure model of wild-type SPF30 were predicted by Alphafold2. A kink-like structure of SPF30 in the C-terminal intrinsically disordered region is colored magenta. (B) Six 293FTR cell lines capable of inducibly expressing deletion mutants (WT, ΔN, ΔTudor, ΔC1, ΔC4, and ΔC5) of SPF30 were treated without (-) or with (+) Dox for 24 h, and FLAG-tagged SPF30 were detected using anti-FLAG antibody, respectively. GAPDH was shown as loading control. 293FTR cell line was used as a parental cell. (C) The recombinants of 6xHis-tagged SPF30 deletion mutants (ΔC1-ΔC6) were purified using Ni-NTA agarose from E. coli strain, and confirmed by SDS-PAGE and CBB stanning. Indicated amounts of proteins were loaded to SDS-PAGE. (D) The structure model of C-terminal α-helix and intrinsically disordered region is shown. A kink-like structure of SPF30 in the C-terminal intrinsically disordered region is colored magenta.

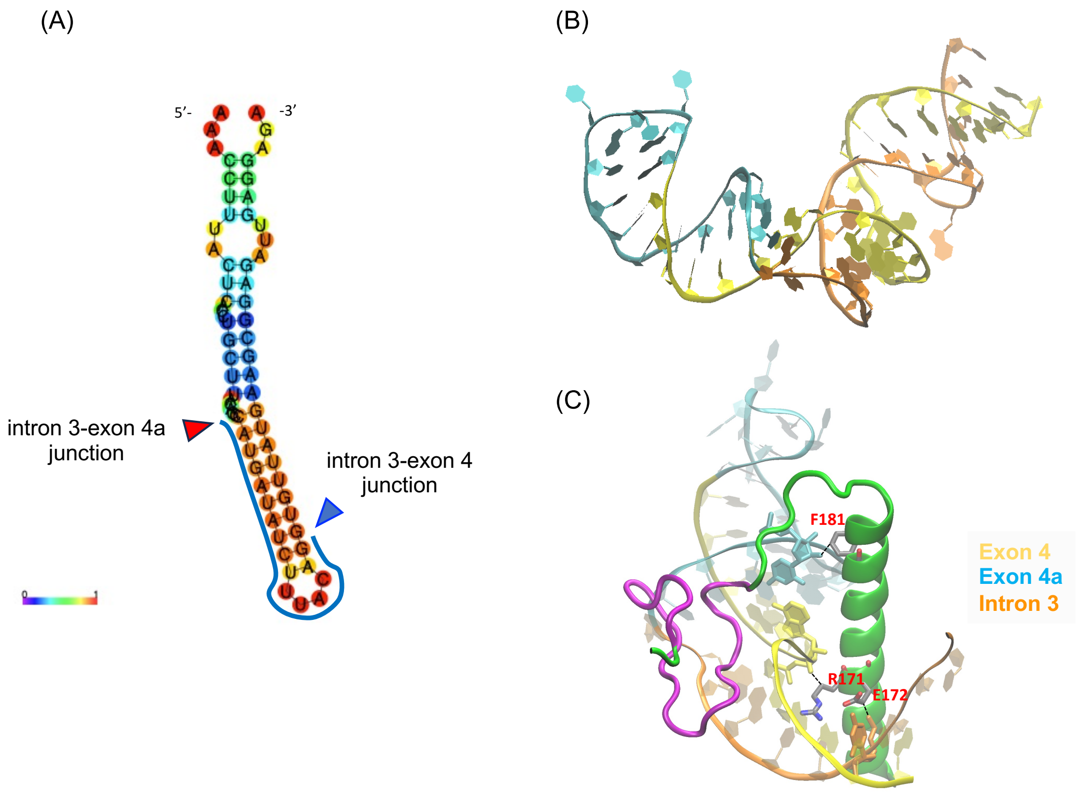


**Supplementary Figure S5.**

The secondary structure model of SPF30 transcript with exon 4a, predicted using the Vienna RNAfold (A) and the RNAcomposer (B). (A) The junction of intron 3 and exon 4a is indicated by a red arrow head, and the junction of intron 3 and exon 4 is by a blue arrow head. The blue line shows the 17 bp unique sequence of exon 4a. (B) Exon 4 is colored yellow, the unique sequence of exon 4a is cyan, and intron 3 is orange. (C) View of the C-terminal α-helix of SPF30 bound to RNA.

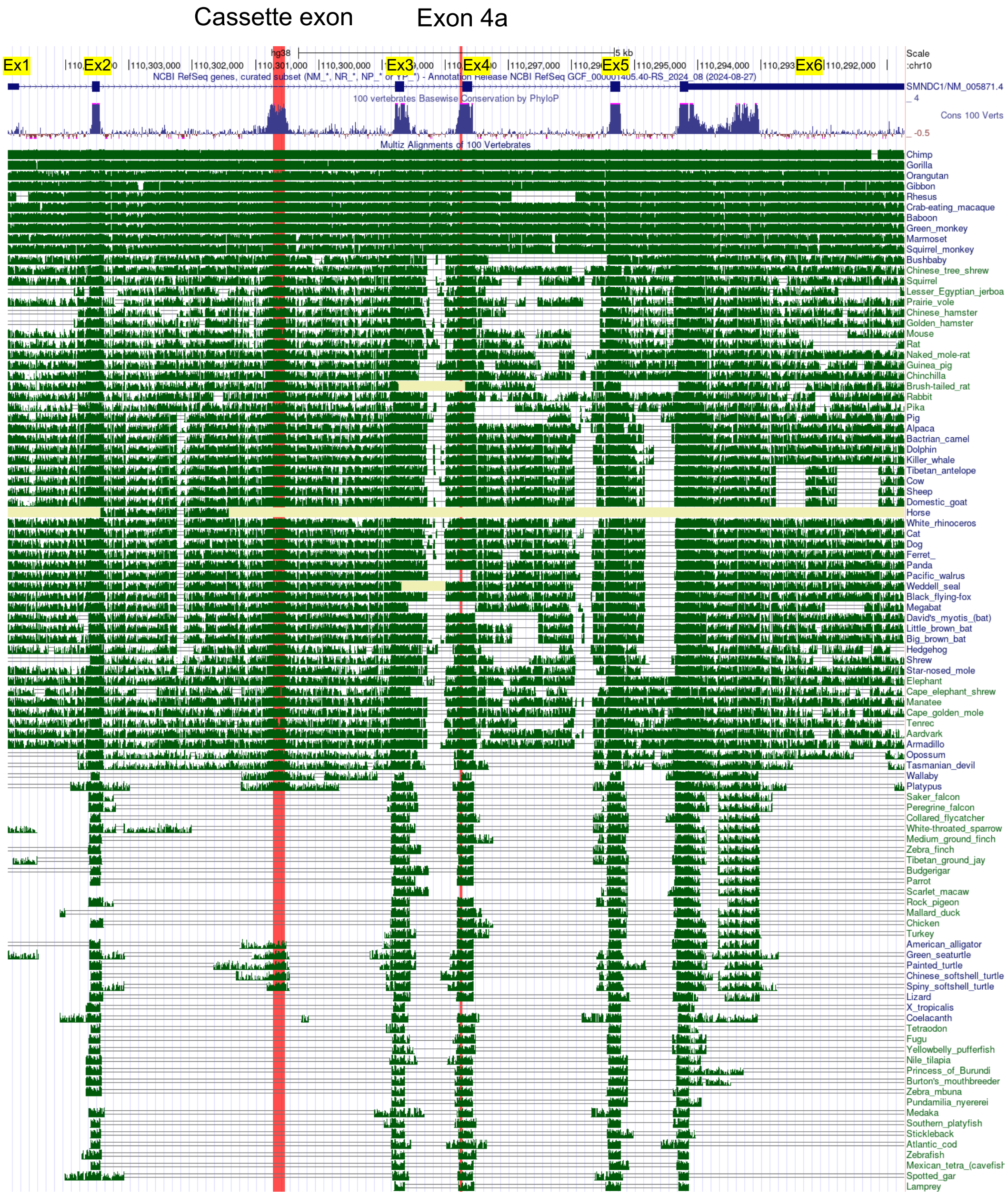


**Supplementary Figure S6.**

Comparison of conserved sequence of SPF30 gene (*SMNDC1*) among vertebrate species. The positions of exon 4a and the cassette exon within intron 2 are shown red lines. As a reference, human *SMNDC1* mRNA and DNA (NM_005871 and GRCh38/hg38, ch10) is used.

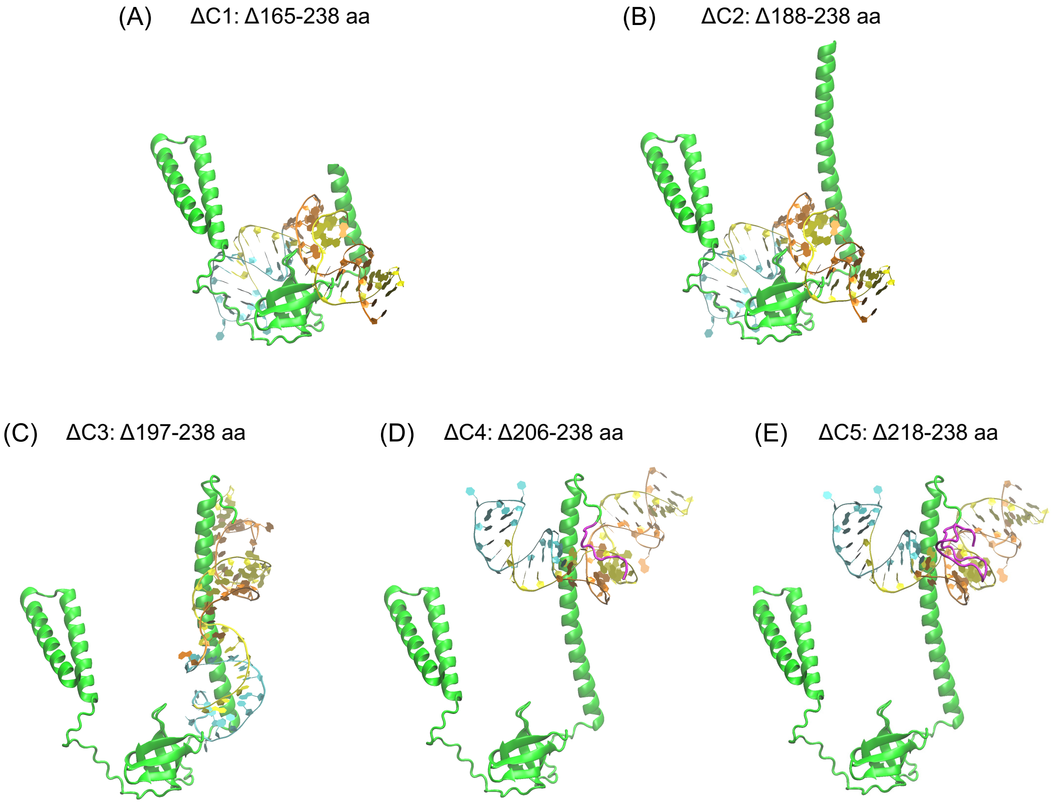


**Supplementary Figure S7.**

Docking models of SPF30 deletion mutants (A; ΔC1, B; ΔC2, C; ΔC3, D; ΔC4, E; ΔC5) and SPF30 transcripts containing exon 4a were predicted by HDOCK software. The kink-like structure of SPF30 is colored magenta. Exon 4 is colored yellow, the unique sequence of exon 4a is cyan, and intron 3 is orange.
